# Multi-team conflict resolution is ineffective for stable decision making

**DOI:** 10.1101/2025.10.06.679762

**Authors:** Kushal Haldar, Vaibhav Anand, Aditya Moger, Tomas Gedeon, Kishore Hari, Mohit Kumar Jolly

## Abstract

Competition between distinct groups is fundamental to complex systems, from political coalitions and economic markets to gene regulatory networks (GRNs) controlling cell fate. Strikingly, cell-fate decision systems predominantly employ binary competition between two mutually inhibitory teams of genes, despite multi-team competition being common in social and ecological contexts. This raises a fundamental question: does binary decision-making offer functional advantages over multi-team systems? We systematically analyzed multi-team competitive networks using Boolean dynamics on signed directed graphs, extending well-characterized two-team GRN architectures to systems with three or more mutually inhibitory teams. Three-team networks produced significantly less stable outcomes than two-team systems, with steady states showing higher structural frustration and increased sensitivity to perturbations. Multi-team systems exhibited unpredictable transitions even under controlled perturbations, especially at lower densities typical of real networks. Networks with four or more teams failed to maintain distinct stable states entirely. Using spectral analysis, we show that team structure can be predicted from network eigenproperties, extending structural balance theory to directed signed networks. Our findings explain why binary decisions dominate biology and provide insights into coalition instability in social systems, market dynamics, and organizational structures. This work establishes fundamental stability principles for competitive networks across disciplines.

## 1. Introduction

Competition is fundamental to many complex systems that span the social, political, economic, ecological, and biological domains. In politics, parties compete for power. In wars, armies vie for resources. In economics, firms compete for market dominance, while in sports, teams or athletes vie for supremacy. In ecological systems, microbial species compete for resources within shared ecosystems [1]. Similarly, in biological systems, competing transcription factors give rise to cell fate decision systems, allowing cells to adopt different phenotypes [2]. These networks of mutually inhibiting transcription factors are found in various contexts, from development (e.g., PU.1-GATA1 mutual inhibition in hematopoietic stem cell differentiation [3]) to disease (e.g., MITF-AXL mutual inhibition in melanoma, controlling the proliferative-invasive phenotype switch [4]). The strength (population/votes/expression level) of the competing entities evolves over time, and depending upon various factors such as the initial configuration, number of competing parties, and the extent of competition, one or more entities can prevail at the steady state. The fate or phenotype of the system is thus determined by the prevailing entities at steady state.

A striking feature of competing systems is the prevalence of binary competition, where only two groups compete at a time. For instance, the cascade of differentiations in embryonic development is driven by a series of binary decisions [5, 6]. Such binary decisions, found in multiple biological contexts such as proliferative vs. non-proliferative transition in melanoma [7], neuroendocrine vs. non-neuroendocrine transition in small-cell lung cancer (SCLC) [8], and gonadal cell-fate transitions during development [9, 10], are results of regulatory networks formed by two competing teams of genes. The interactions within a team are positive and interactions across teams are negative, ensuring two dominant phenotypes at steady state, each characterized by the coexpression of nodes from only one of the teams [10]. Additionally, these networks allow for transition between these phenotypes upon perturbation. Multi-fate systems, such as T-cell differentiation [11], that involve multiple competing teams are rarely observed in biological regulatory networks. This raises the questions: Is binary decision-making functionally advantageous over multi-fate systems? How do multi-team systems differ dynamically from two-team systems? What features characterize multi-team competition in GRNs or other competitive networks?

To address these questions, we compared the dynamic behaviors of binary GRNs with multi-fate GRNs. Inspired by the network structures of binary cell-fate decision systems [12] and T-cell differentiation network [11], we constructed artificial networks of two and three teams that mutually inhibit each other. We generated large ensembles of such networks with varying densities and team sizes. We then simulated these networks using a threshold-based Boolean formalism for a large number of initial conditions, and calculated the frequency distributions of the steady states over these initial conditions. We analyzed these networks to compare the stability of “single-positive” phenotypes that represent terminally differentiated/functionally modular cellular phenotypes and transition dynamics between different steady states under perturbations. Our results demonstrate that three-team networks produce states that are less stable and exhibit more unpredictable transitions compared to two-team networks. Extending this analysis to networks with more than three teams, we found that such networks fail to maintain distinct terminally differentiated states with unique expression patterns at a biologically reasonable network density.

Reliable transitions are crucial for maintaining proper cellular functions in biology. Unreliable transitions contribute to diseases like cancer, where uncontrolled or misdirected differentiation promotes metastasis. In regenerative processes, precise and stable transitions are necessary for proper tissue regeneration. Similarly, in social systems, unstable transitions can lead to political volatility or economic disruptions [13]. Because robustness and reliable transitions are critical across these systems, our findings suggest that binary decision systems offer a functional advantage over multi-fate systems by ensuring more stable and predictable outcomes.

## 2. Results

### 2.1. Stability of the steady states of two vs three team networks

Teams of elements/nodes are commonly found in many contexts of social and biological networks. Networks with a single team of nodes, and those with two mutually inhibiting teams have the property of structural balance, in that the number of inhibitions involved in three connected nodes is always even [14]. Generally speaking, any feedback loop in an ideal two-team network is positive. While more than two mutually inhibiting teams are seen in social contexts [13], they are rarely observed in decision-making at a cellular level. We hypothesize that the reason for the difference in abundance can lie in the difference in the dynamics of two vs multiple team networks. To evaluate the dynamics, we chose to simulate artificially generated networks with teams using a threshold-based Boolean formalism. Threshold-based rules are also commonly employed in social networks to study their dynamics. The Boolean formalism is inspired by the Ising model used to model ferromagnetic material and spin glasses [15]. Briefly, the formalism assumes that the expression level/state of each node in the network can be either +1 (high expression) or -1 (low). The state of the network *S*(*t*), hence, is a vector of N (= number of nodes) binary variables. Using the update rules described in Eq. 3, we simulate each network starting from 100,000 randomly chosen initial states until a steady state is reached. The steady states of two-team networks are primarily composed of all nodes from one team having the state +1 and all the nodes in the other team having the state -1 **(Fig. 1A)**. We refer to such states as single-team positive or single positive, in short.

**Figure 1.**
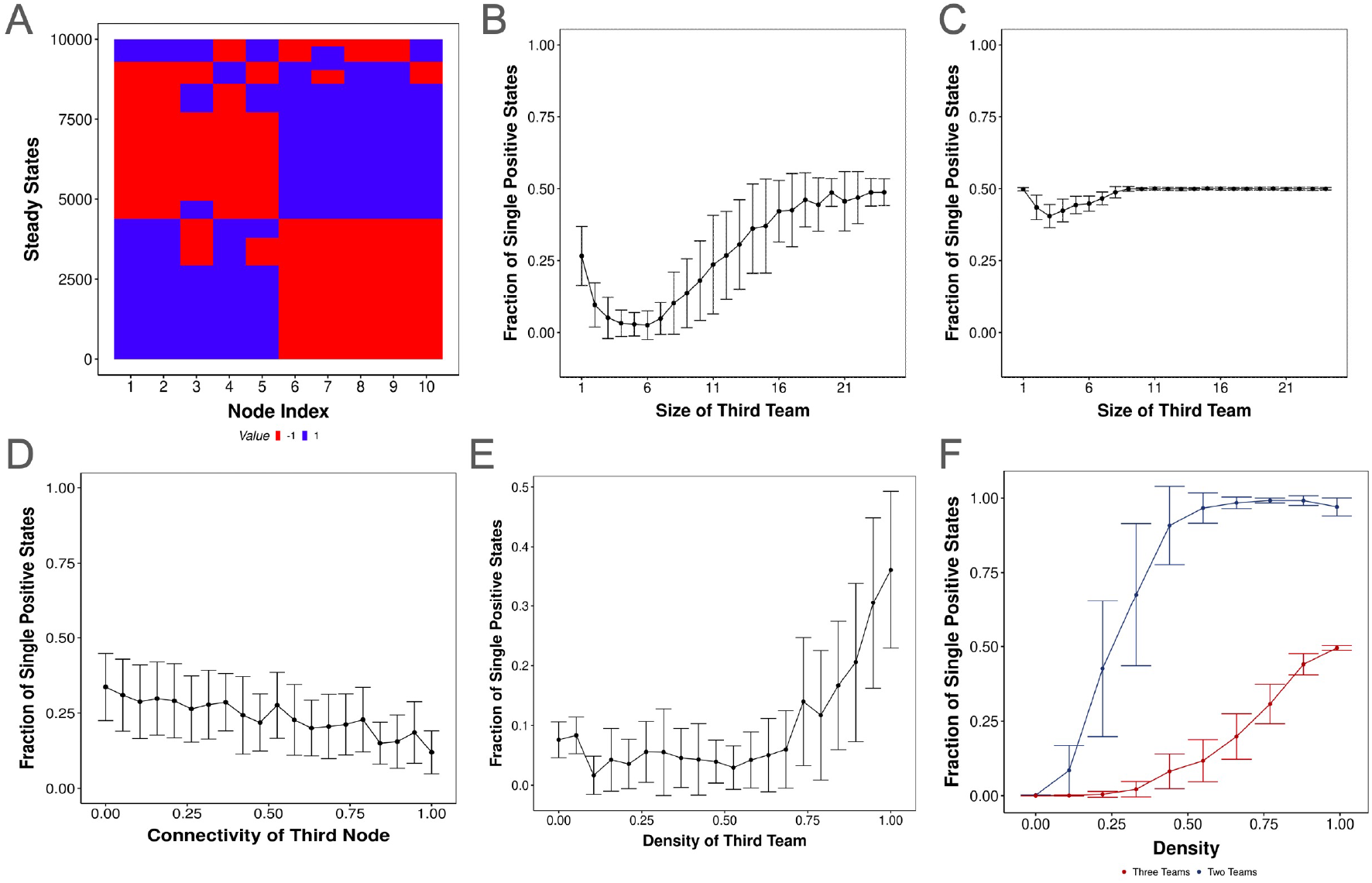
Three team networks lead to a reduced incidence of “pure” states. **A)** Heatmap shows the steady states of a two-team network with 0.3 density. Each cell corresponds to the expression level of the corresponding node (column) in the corresponding steady state (row). Red is a high expression level, and blue is low. **B-C)** Line plot depicting the mean ± sd fraction of single positive steady states over multiple networks of two teams with 5 nodes per team, and a third team of varying number of nodes from 1 to 24. The edge density is **(B)** 0.3 and **(C)** 0.9. **D)** Mean ± sd of the fraction of single positive states as a function of the degree of an external node inhibiting all other nodes in a two-team network, thereby serving as a third “team”. **E)** Mean ± sd of the fraction of single positive states as a function of the density of a third team inhibiting all other nodes in a two-team network.

A two-team network can have two single positive states, one for each team, and we see that both the states occur with a similar frequency, suggesting that the network allows for either team to “win” with equal probability. A small fraction of initial conditions also converge to steady states that are not single positive. We refer to these states as hybrid states because a subset of nodes from both teams have +1 expression, unlike the single positive states. While *>* 90% of the initial conditions converge to a single positive state, a very small fraction of initial conditions converge to the hybrid states. At a higher edge density (*E/N* ^2^ where *E* is the number of edges and *N* is the number of nodes in the network), all initial conditions converge to single positive states **(Fig S1A, i)**

To understand the dynamics of multi-team networks, we started by analyzing three team networks. Upon the two-team network, we started to build a third team by adding a set of nodes that inhibit and are inhibited by both teams (team A and team B) currently in the network. We generated 50 such networks for each size of the third team (team C), ranging from 1 to 24, with a constant edge density of 0.3, and obtained the corresponding steady states in the same manner as described before. We find that the fraction of initial conditions converging to any single positive state has now decreased significantly, from *>* 90% for a two-team network to 25% with a single node in team C. As we further increased the size of team C, the fraction of single positive states further reduced, approaching 0% for a team C of size 5. The steady states did not seem to follow any particular configuration as they did in two-team networks at the same density **(Fig S1A, ii)**. When the size of team C was equal to that of A and B (5 nodes), almost all of the steady states were hybrid for three-team networks at 0.3 density. To determine if three-team networks are capable of any structured steady state configuration at all, we repeated the same experiment at an increased density of 0.9. At higher density, the fraction of single positive states increased to 50% when the size of team C was 1. We did observe a decrease in the fraction as the team C size increased, but the magnitude of the decrease was not as significant as that at 0.3 density. Furthermore, the minimum fraction of single positive states (40%) was observed when the size of team C was 3, after which the fraction started to increase. Looking at the steady state configurations, we identified that in addition to single positive states, a significant fraction of the state space converged to steady states where nodes from two of the three teams have +1 state and the nodes from the remaining team have-1 state. We refer to these states as double positive states. Note that the double positive states are mirror states of single positive states. If one takes a single positive state and flips the state of all the nodes (from -1 to +1 and vice versa), it would turn into a double positive state. In two-team networks, the two single positive states are mirror states. Mirror states have the same frustration value, and due to the symmetry of the formalism, the frequency of the fraction of double positive states also mirrors that of single positive states, both at low and high density **(Fig S1B, C)**.

All the states that could not be classified as single positive or double positive states are referred to as hybrid states. Correspondingly, as the fraction of single and double positive states decreases with increasing number of nodes in team C, the fraction of hybrid states increases. As the number of nodes in team C increase further, the fraction of single positive and double positive states increased for both densities, saturating at a mean of 50% each. To understand this increase, we took a closer look at the composition of the different single and double positive states observed at different sizes of team C **(Fig S2A-B)**. At a team size of 1 for team C, nearly all of the single positive states have the configuration of Abc (all nodes of team A have a state +1 and all the other nodes have a state -1) and aBc. As the size of team C increases, the fraction of the abC state increases, while that of Abc and aBc decreases, contributing to the initial decrease. At larger sizes of team C, team C completely dominates teams A and B, thereby making abC the dominant single positive state. Overall, we find that an imbalance in team sizes leads to the state space orienting towards the larger team.

We further explored the effect of the imbalance by varying the density, i.e., the ratio of the edges present and the total number of possible edges, within team C and between team C and the other two teams, while keeping the size of team C constant, and the density of the edges within and between teams A and B at 0.3. When team C has a single node, the density variation represents the variation in the connectivity of team C to teams A and B **(Fig 1D)**. We find a steady decrease in the total fraction of single positive states as the connectivity of team C increases. However, for a team C size of 3 and higher, where we could manipulate the density within team C as well, the fraction of single positive states increases with increasing density **(Fig 1E)**. A closer look at the composition of the single positive states reveals that the increase in single positive state frequency is due to an increase in the abC states for larger size of team C when there are intra team interactions in addition to inter-team interactions, a feature lacking in single-node team C (**Fig S2C-E**), suggesting that a team of nodes has better unity than a single node. We further simulated balanced, symmetric networks with two and three teams at different densities and found that not only do the two-team networks have a higher fraction of single positive states at all values of density, but the fraction saturates quickly than that of three teams.

Frustration of a steady state is a measure of the extent of mismatch between the network structure and the expression levels of the nodes. An edge is said to be frustrated if the product of the states of the nodes involved in the edge and the sign of the edge is -1 (*J*_*i*_*js*_*i*_*s*_*j*_ = −1). Steady states with high frustration (a large fraction of frustrated edges) are more sensitive to perturbations and are therefore less stable [10, 16]. We compared the frustration of the steady states obtained from two-team networks vs three-team networks at 0.3 density. While the minimum value of frustration for two team networks was 0, the same for the three team networks was 0.2, suggesting that at least 20% of the edges are always frustrated at steady state for three team networks **(Fig 2A)**. The average frustration of the three-team steady states is much higher than the team steady states. We simulated 50 networks, each of two and three teams with a density of 0.3, and calculated the mean frustration for each network across all steady states of the network. The highest mean frustration across 50 two-team networks was lower than the lowest mean frustration across 50 three-team networks, suggesting that the steady states emergent from two teams are more resilient to perturbations than those from three teams. Furthermore, the mean frustration of three team networks has lower variance **(Fig 2B)** than two team networks. This difference in variance is maintained for steady states within a single network as well (**Fig 2A**). This difference in variance highlights the fundamental difference in feedback within two and three-team network topologies. While two-team networks only have positive feedback loops, three-team networks have an abundance of negative feedback loops. Consider the negative feedback loop between two nodes: X inhibiting Y while Y activating X. In such a loop, one of the two edges is always frustrated, no matter what the configuration of states. This property of negative feedback loops ensures a high mean frustration for three-team networks. Two-team networks, on the other hand, do not have negative feedback loops, allowing the existence of low frustration states. Thus, we get dependent on the connectivity of the network; two team networks can have high and low mean frustration.

**Figure 2.**
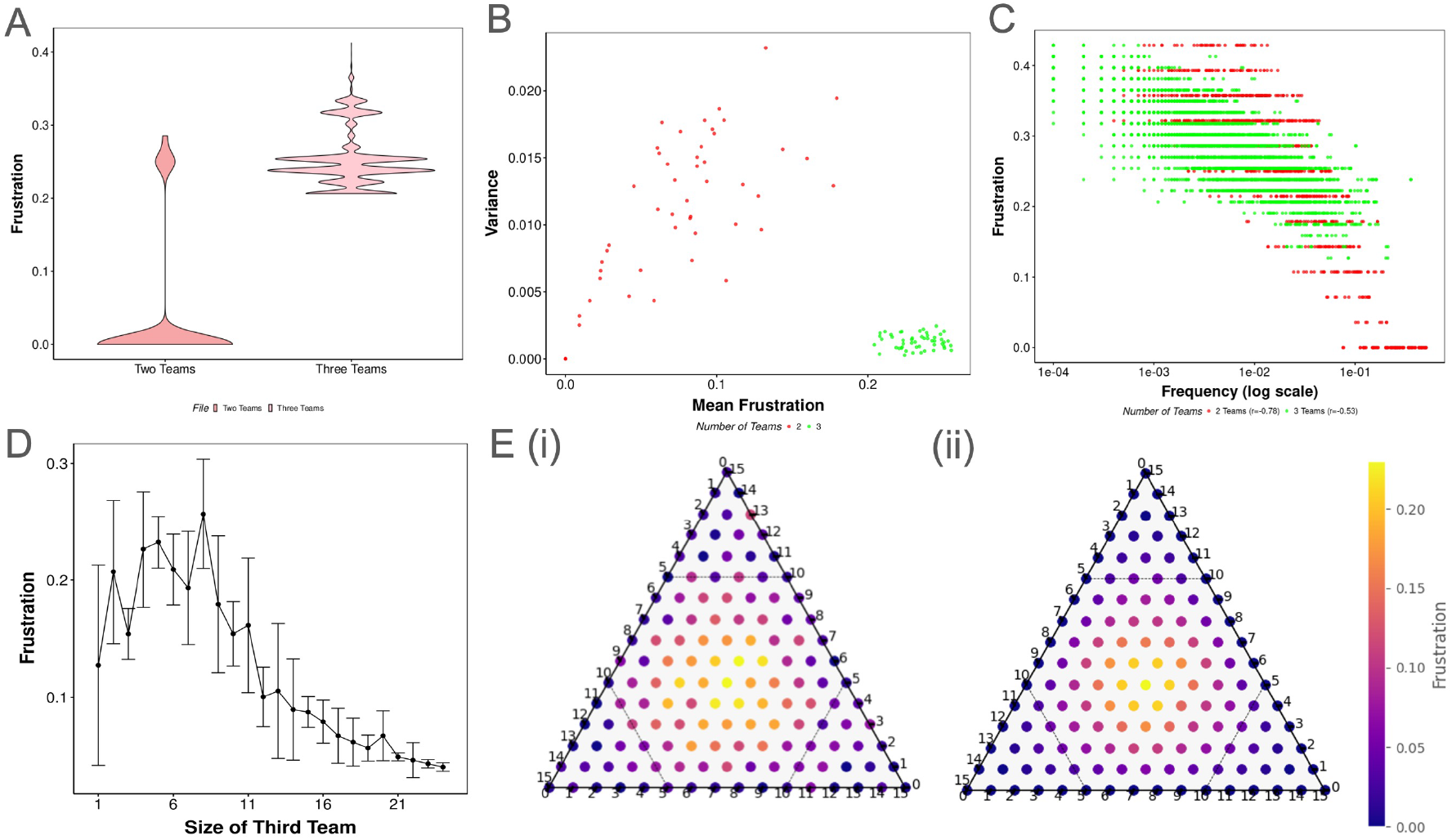
Three-team networks lead to higher frustration in steady states than two-team networks. **A)** Violin plots representing the distribution of the frustration of steady states of one representative network, each with two and three team networks of density 0.3. **B)** A scatter plot depicting the mean and variance of the frustration of steady states for 50 two-team networks (red) and 50 three-team networks (green), each with a density of 0.3. Each point is one network. **C)** A scatter plot of the steady state frequency vs the frustration of the steady states of 50 two (red) and three (green) team networks of 0.3 density. Each point is a single steady state **D)** Line plot describing the mean ± sd frustration of networks of two teams (5 nodes per team) with a third team of varying size. For each point on the x-axis, we sampled 50 networks and pooled the frustration of all steady states of all networks. **E)** Simplex heat maps indicating the frustration of different size configurations of three teams with density (i) 0.3 and (ii) 0.9. Each point represents the mean frustration of 50 networks with three teams, with the size configuration indicated by the sides of the simplex plot. Blue color indicates less frustration, while red/yellow color indicates high frustration.

In both three and two-team networks, frustration of a steady state has a strong negative correlation with its frequency (the fraction of initial conditions that converge to the steady state **Fig 2C**). The strength of the correlation is higher for two-team networks than for three-team networks. When an imbalance in team sizes is introduced into these networks, frustration has the opposite trend to that of the fraction of single positive states. At a density of 0.3, as the size of team C increases from 1 to 5, the mean frustration increases. For team C of 6 nodes and above, we observe a decrease in the mean frustration, approaching zero at large team C (**Fig 2D**). We can attribute this result to the sheer size of team C in comparison to the other two, as most of the initial conditions converge to abC and ABc states. In these states, the frustrated edges are the ones between team A and B, and the number of these edges is much smaller (5 * 5 * 0.3) than the total number of edges, making the frustration 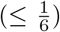 negligible. This reduction in frustration due to imbalance is seen to be independent of density **(Fig 2D)**. We then asked if network frustration changes with the density of three-team networks. We calculated the mean frustration of three team networks with varying team sizes (1-15 nodes) for 0.3 and 0.9 densities **(Fig 2E)**. Interestingly, unlike the fraction of single positive states, the changes in frustration are not dependent on density, as shown by the striking similarity between **Fig 2E i and ii**. Three-team networks reach high frustration only when the team sizes are similar (center of the simplexes). When the team sizes are imbalanced, the steady state space is biased towards the imbalance (for example, abundance of abC states when the C team is much bigger than the A and B teams, (**Fig 1B**)). Overall, three teams lead to less stable, more frustrated steady states as compared to two team networks.

### 2.2. Sensitivity of three team steady states to perturbations

Social and biological systems always face perturbations that affect the state of individual nodes, percolate through the networks, and change their state. It is important to be able to predict the effect of different perturbations in the system for various ends, including inhibiting unwanted transitions. We attempted to interrogate the sensitivity of prominent phenotypes emergent from two-team and three-team networks to various types - simultaneous and sequential - (see corresponding sections in Methods), and levels of perturbations.

We first investigated the effect of simultaneous perturbations, where the states of multiple nodes in a steady state are perturbed simultaneously and allowed to evolve to a steady state, by analysing the evolution of single-positive and double-positive phenotypes emergent from three-team networks in the presence of simultaneous perturbation. We generated an ensemble of 100 networks of three teams with 5 nodes per team and a density of 0.9. We first simulated these networks to obtain the steady states. As shown earlier, these states can be classified into three types: single positive (labelled as ‘+’), double positive (labelled as ‘++’), and hybrid states. For each single positive and double positive state on each network, we generated 100 X N perturbation experiments, where N is the number of nodes in the network. Each experiment consists of simultaneously perturbing *i* ∈ [1, 2, 3…*n*] randomly chosen nodes in the corresponding steady state *S*_0_, simulating the resultant state *S*_1_ using Eq 3 till a steady state 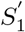, referred to as the destination, is reached. We subsequently obtained the relative frequencies of each destination over all repetitions for each single/double positive state on each network. At low level perturbations (*<*30% nodes) majority of transitions from single positive states were back to the same state, that is, no transition occurs. As perturbation strength increased into the lower-medium range (30-50% nodes), the dominant destination shifted towards double positive phenotypes that were not the mirror states (all nodes having the opposite expression as the original state) of the initial single positive state (for example, if the initial state is Abc, mirror state is aBC, and the non-mirror double positive states are AbC and ABc), with the frequency of this destination type peaking at a perturbation strength of around 40% of the nodes in the network. Further increase in perturbation strength into the higher-medium range (50-70%) led to a decline in the frequency of transitions to non-mirror double positive states and a corresponding increase in frequency of single positive states not identical to the initial single positive state. This destination type had a maximum frequency at a perturbation level of around 60% of the nodes in the network; further increase in perturbation led to a decline in frequency of single-positive destinations and a rise in the frequency of the double positive destination, mirroring the initial single-positive state **(Fig 3A)**. This pattern is intuitive considering the hamming distances of the respective destination types from the initial state. For example, among the available steady states for three-team networks with a density of 0.9, the immediate neighbours of a single positive state are two double positive states that have an additional team on. An analogous pattern is observed for double positive initial states, where the dominant phenotype shifts from the same state, to non-mirror single positive states, to other double positive states, and finally to the mirroring single-positive state as the perturbation level increases from 0 to 1 **(Fig 3B)**. Each network also possesses a smaller fraction of relatively less stable hybrid steady states, due to which, at any level of perturbation, there is a small probability of reaching a hybrid destination that is neither single nor double positive. At a density of 0.3, most of the steady states are hybrid **(Fig 1B, S1A, ii)**. Thus, transitions starting from single positive and double positive states also end up in hybrid states. Given the number of unique hybrid states, it becomes hard to predict the destination of state transitions at lower densities in three teams. However, as shown in **Fig 1F**, two teams also have a significantly lower fraction of single positive steady states and therefore a higher fraction of hybrid states at lower density values. Thus, we wanted to directly compare the sensitivity of single positive states to perturbations in both two and three-team networks.

**Figure 3.**
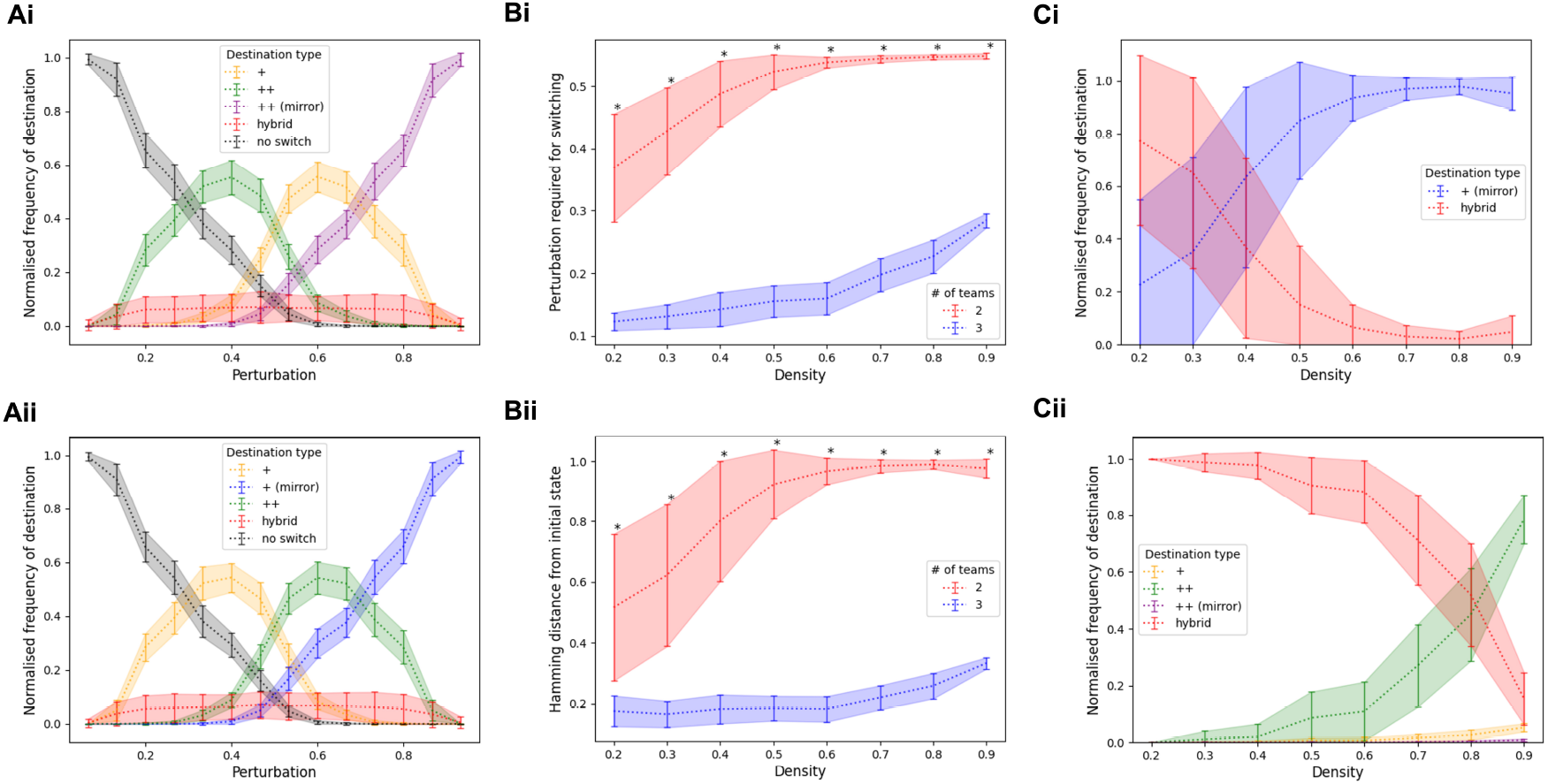
Patterns of state transition in two and three team networks Ai. Line plot depicting the fraction of different destinations that single positive states get to after a given extent of perturbation. The X-axis represents the fraction of nodes being perturbed (flipped signs). Different destinations are represented by different colors. Network density=0.9. **Aii** - Line plot depicting the fraction of different destinations that double positive states get to after a given extent of perturbation. Network density=0.9. **Bi** - Line plot depicting the amount of sequential perturbation required to transition out of a single positive state at a given density. The Y-axis denotes the fraction of nodes perturbed. Different numbers of teams are represented by different colors. **Bii** - Line plot depicting the hamming distance of destinations after switching out of a single positive state under sequential perturbation at a given density. **Ci**- Line plot depicting the fraction of different destinations that single positive states get to after sequential perturbation at a given density in networks with 2 teams. **Cii**- Line plot depicting the fraction of different destinations that single positive states get to after sequential perturbation at a given density in networks with 3 teams.

We subsequently investigated the effect of sequential perturbation, where nodes in a steady state are perturbed one at a time till the system evolves to a new steady state distinct from the initial state. For each single positive steady state in our ensemble of networks, we perform the sequential perturbation experiment and asked what is the extent of perturbation required to switch the state **(Fig 3B, i)** and the hamming distance of the perturbed state from the single positive state **(Fig 3B, ii)**. At every density value, we clearly see that the single positive states from three team networks are much more sensitive to perturbations than those from two team networks. At lower densities, perturbing less than 15% nodes (2 nodes) is enough to switch the state. On the other hand, even at the lowest density, single positive states of two-team networks were resistant to a perturbation of up to 40% nodes on average before switching. Interestingly, the mean hamming distance of the switched state for three teams at lower densities is also around 0.15. Note that the hamming distance from a single positive state to the nearest double positive states is 1/3 and the nearest single positive states is 2/3, suggesting that the transition at lower densities is to hybrid states. Single positive states of two-team networks also transition to hybrid states, as the hamming distance between two single positive states in two-team networks is 1, and the average hamming distance of the destination is 0.5. As the density increases, the perturbation required to switch the state increases in both two and three teams, but the rate of increase is much higher for two-team networks. At high density values, single positive states in two team networks are resilient up to 50% of nodes perturbed, after which the state switches to the mirror state (hamming distance 1). Three team networks, on the other hand, lead to a switch in the state after perturbation in 25% of the nodes. Furthermore, while two teams see a high resilience and near perfect switch from single positive to single positive state from a density of 0.6 **(Fig 3C i)**, three team networks maintain high sensitivity and switch to hybrid states until 0.6 density, and reach the next double positive state (hamming distance 1/3) only at 0.9 density **(Fig 3C ii)**. Overall, while both two-team and three-team networks require high density for the state transitions when exposed to random perturbations to be predictable (i.e., complete transitions are achieved in the direction of perturbation), the predictability (probability of successful transitions) is much lower in three-team networks, and only achieved at very high density values.

### 2.3. Destination-driven perturbations reveal the unreliability of three team networks

Our analysis so far has given us a reasonable understanding of the phenotypic landscapes of two-team and three-team networks. We have also investigated the behavior of the steady states in the presence of different levels of randomized perturbations and expected transitions of single and double positive states. We now ask if we can use this knowledge to try and control the direction in which the state transition happens. Particularly, we are interested in transitions from single positive to single positive states, since these states represent meaningful phenotypes such as cell fates. Particularly, we seek the answer to three questions: 1) The set of nodes that should be perturbed to achieve the transition, 2) The number of nodes that need to be perturbed to achieve the transition, and 3) Probability of success given the perturbation in the right direction.

For two-team networks, the single positive states are mirror states; therefore, the superset of nodes to be perturbed constitutes all nodes. At a higher density of the network, we saw in **Fig 3B** that the average perturbation required for a successful transition from single positive to single positive state is 0.55. To understand which nodes must be perturbed, we generated a two-team network with 0.9 density, and repeatedly perturbed a single positive state with combinations of node perturbations multiple times each **(Fig 4A)**. We find that perturbing 6/10 nodes is guaranteed to cause a transition, but when 5 nodes are perturbed, the probability of a successful transition is 0.5 **(Fig 4A)**. Due to the symmetry of the network, any combination of 6 nodes (i.e., 3 from each team or 4 from one team, two from the other, etc.) can initiate a successful transition from single positive to single positive states at high density.

**Figure 4.**
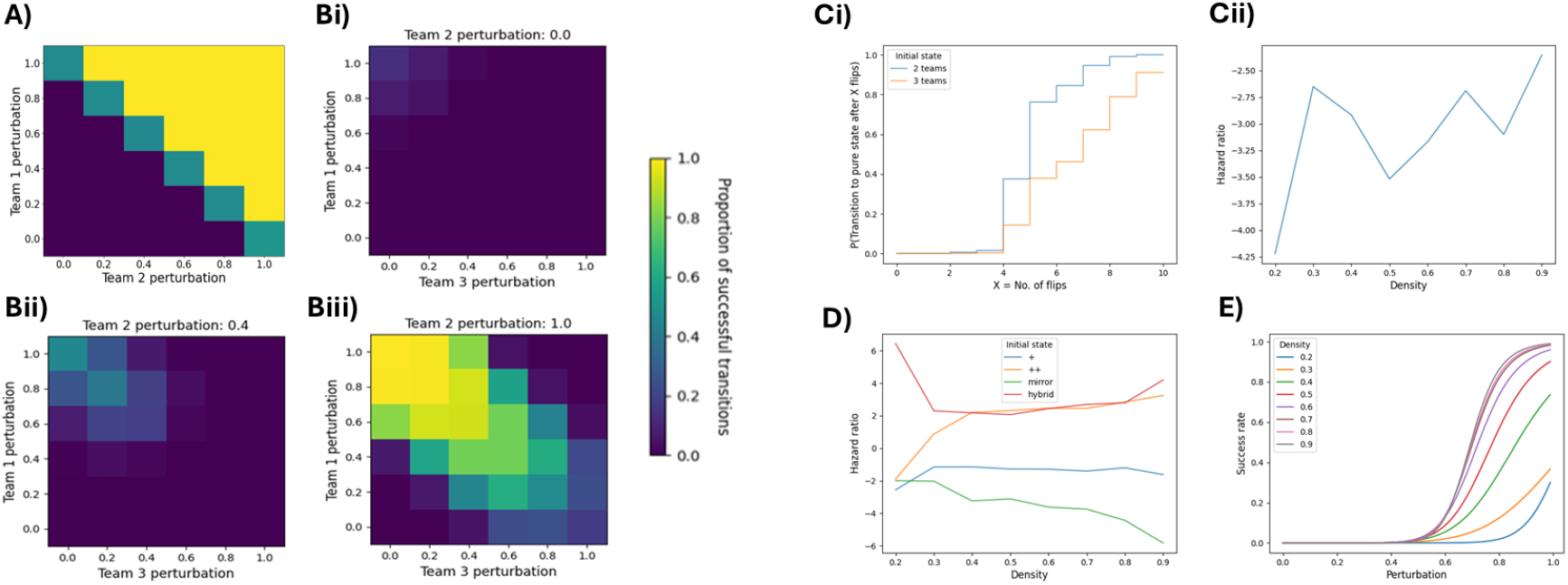
Dynamics of state transition in two and three team networks. **A** Heatmap depicting the success rate of transitioning from state 10 to state 01 on perturbing different proportions of each team in a two-team network with 0.9 density. Each axis represents the proportion of nodes that have been flipped in the corresponding team **B**. Heatmap depicting the success rate of transitioning from 100 to 010 on perturbing different proportions of each team in three-team networks with 0.9 density. **Ci** Step plot depicting the probability of a successful transition from a single positive state to a different single positive state at a given level of targeted sequential perturbation in two-team and three-team networks at 0.9 density. **Cii** Hazard ratio of successful transition of three team networks as compared to two team networks for different densities. **D** Line plot depicting hazard ratios of different types of initial states for transitioning to single positive final states under targeted sequential perturbation at different densities in three-team networks. **E** Success rate for transitions between single positive phenotypes at a given level of targeted perturbation in three-team networks. Different network densities are represented by different colors.

The transitions in the three teams get more complicated. We present an example of transition from Abc to aBc state in **Fig 4B** for a three-team network of 0.9 density, repeating the same experiment as with the two-team network above. For achieving the desired transition, the perturbations must ensure that team A is turned off, team B is turned on, and the expression of team C is not changed. Thus, the set of nodes that can guarantee Abc to aBc transition is the 10 nodes from teams A and B. For a single positive to single positive transition in two teams, while perturbing all 10 nodes guarantees the perturbation, turning off the active team or turning on the inactive team (i.e., perturbing half the nodes) was enough to cause a successful transition in at least 50% of cases. However, the fraction was much lower in three-team networks, even at high density (perturbing team A **Fig 4B, i**). As we start to perturb team B, we find the fraction of cases with successful transition increasing to around 0.4 **(Fig 4B, ii)**. It was also not enough to just turn the team B on to ensure a successful transition. This behavior is expected, since the perturbation leads to the double positive ABc state where it gets stabilized **(Fig 3B, iii)**. A successful transition in all cases is achieved only when we perturbed all nodes of team B and *>*60% of the nodes in team A. Therefore, the minimum perturbation required in the direction of the transition is 9/10 nodes, a much higher number than the 6/10 needed for two teams. It is further important to note that the proportion of successful transitions decreases as we perturb team C beyond 20%. Similar observations were made for other state transitions for three team networks where *>* 80% nodes needed to be perturbed for successful transitions **(Fig S4)**.

We present a comparison of two vs three teams by calculating the probability of transition from single positive to single positive states across multiple networks of 0.9 density, given X nodes perturbed in the desired direction **(Fig 4C, i)**. Below 4 flips, we do not achieve any transition in both cases, but above 4 flips, the team network always outperforms the three-team network by having a higher probability of transition to a single positive state. Interestingly, at 10 nodes, the probability of successful transitions is 0.9 for three teams compared to 1 of two teams, indicating that even when exposing the single positive state to the maximum extent of perturbation possible at high density, a small fraction of uncertainty remains in three-team networks. We fit this data generated across all density values to the Cox proportional hazard model and found, unsurprisingly, that two teams always have a better probability of successful transition as compared to three teams **(Fig 4C, ii**, hazard ratio *>* 1).

We further investigated the rate of successful transitions from other types of states in three team networks to the closest single positive states. The probability of success had the inverse order as the relative hamming distance from the closest single positive state, with highest probability achieved for hybrid states and the lowest for the mirror state across multiple density values **(Fig S5A)**. We then fit the combined directed state transition data of two and three teams to the Cox model to compare the relative success rate of transition to single positive states starting from different initial states **(Fig 4D)**. At low density, the transition probability was low for all initial conditions except for hybrid states. As density increases, hybrid and double positive initial states end up having similar hazard ratios, greater than those of single positive and mirror states.

Finally, we asked how successful one can be in controlling the transitions starting from a single positive steady state to any other desired state, whether it be another single positive state or a double positive state. We know in two-team networks that the transition to another single positive state has a high success rate, and correspondingly, the transition to hybrid states will have a much lower success rate. For three teams, however, the behavior is not as clean. Two-team networks show a much higher success rate for directed transitions as compared to three-team networks, especially at lower density values **(Fig S5B)**. While the success rate does increase monotonically with increasing directed perturbation as expected, the mean success rate even at maximum perturbation does not approach 1 at density values below 0.7 **(Fig 4E, S6A, i)**. Furthermore, the variance in success rate changes non-monotonically with increasing perturbation level, with the maximum values seen at a directed perturbation level of 0.8. At low levels of perturbation, the driving force is not strong enough to get any successful perturbations, while at high perturbation levels, the success rate is consistently higher, explaining the low variance at both ends. At moderate levels of perturbation, the specific perturbation as well as the variation introduced by network topology play a role, increasing the variance of successful transitions. As the signals of cell fate decisions often only affect a relatively smaller fraction of nodes, and because biological networks are sparse, multi-fate decisions driven by three-team networks are highly unreliable for robust cell fate decision systems.

### 2.4. Multiple mutually inhibiting teams are incapable of reliable terminal cell fate decisions

After exploring two and three-team networks for their potential as cell-fate decision systems, we then explored the potential of multi-team networks to regulate multi-fate systems. To our knowledge, the regulatory networks that can allow the co-existence of more than 3 phenotypes with mutually exclusive expression patterns have not been characterized. Achieving the coexistence of multiple single-positive steady states would require each team to be able to suppress the expression of all other teams. Thus, we evaluated networks of multiple teams inhibiting each other as candidates of multi-fate systems. We generated artificial networks having a higher number of teams (4-7) with each team having 5 nodes, using the stochastic block model. We maintained a density of 0.3 to accommodate the sparsity of biological networks. We first observe that only 2 and 3 team networks are capable of resulting non-zero percentage of single positive states, and this property is independent of density **(Fig 5A)**. At high density, n-team networks show a dominant presence of states with n/2 or (n-1)/2 and (n+1)/2 teams on, depending on whether n is even or odd. For example, four and five-team networks at high densities lead to double positive states and double and triple positive states, respectively, at high density [17]. At low density, however, five-team networks do not show any significant frequency of double or triple positive states, while four-team networks show a low, yet significant frequency of double positive states, higher than three-team networks. Six and seven-team networks do not show any such states at low density.

**Figure 5.**
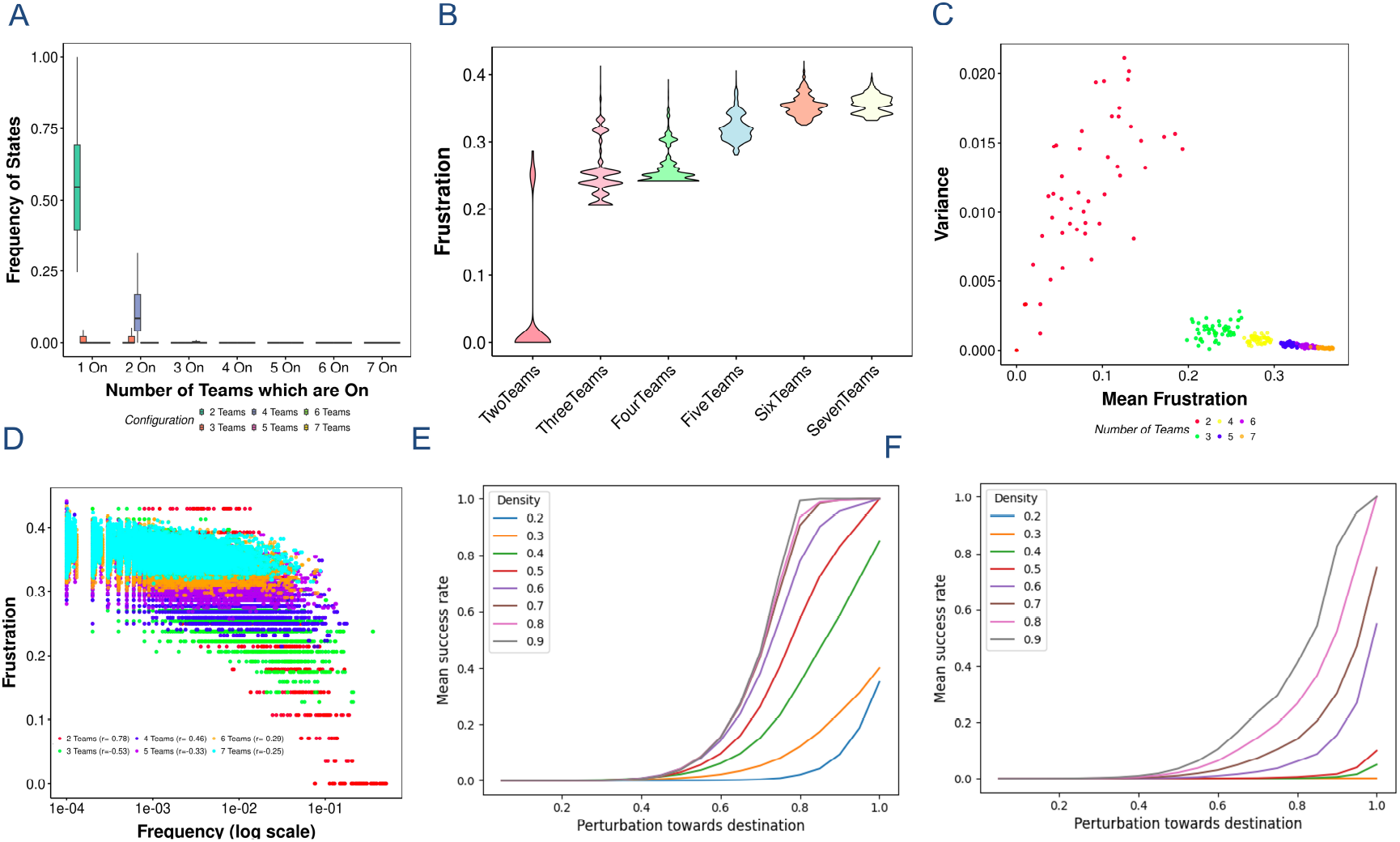
Phenotypic landscape of multiple mutually inhibiting teams of nodes. **A** Frequency of n-positive steady states for networks of multiple mutually inhibiting teams (2-7, colored). **B** Violin plots depicting the distributions of frustration of the steady states of n-team networks. **C** Scatterplot depicting the mean vs variance of frustration of steady states of n-team networks. **D** Frustration vs steady state frequency of steady states from n-team networks. The corresponding correlation values for each n-team network are provided in the legend. **E** Line plot showing the mean success rate of transition from double positive states to other dominant states in four team networks. Different colored lines represent different densities. **F** Same as E for five-team networks

Networks with a higher number of teams continue to have steady states with high frustration as compared to a two-team network. Networks with a higher number of teams also follow the properties exhibited by three-team networks. 1) Representative networks of each type at 0.3 density have steady states which have higher frustration **(Fig 5B)**. 2) The distribution of frustration of steady states has a high mean and a low variance. There is a minor increase in mean frustration as the number of teams increases from three onwards **(Fig 5C)**. 3) The magnitude of negative correlation between the frequency and frustration of the steady states reduces as we increase the number of teams **(Fig 5D)**.

We further performed state transition experiments of networks with four and five teams as we did in **Fig 4E, F**. Four team networks showed a better mean success rate as compared to three teams, with a success rate approaching 1 at densities above 0.4. At a density of 0.4, the maximum transition rate of 0.8 was observed, making the four-team network a much better candidate for robust cell fate transitions as compared to three-team networks. Furthermore, the success rate shows a saturation at about 0.7 perturbation level at higher densities, similar to that of two-team networks. The variance in success rate also increases with the extent of perturbation as before, but maxes out at 0.7, decreasing afterwards **(Fig S6B)**. Five team networks, on the other hand, have similar to worse success rates as compared to three team networks. Overall, we find that a team network is the best candidate for robust cell-fate decision systems, followed by four-team networks.

### 2.5. Spectral properties of teamed networks

So far, we have characterized the landscapes emergent from multi-team systems for their steady states and state transitions. Based on our analysis, it is clear that identifying the number of teams in a given system would provide useful insights into the expected configuration of the steady states as well as the directions and probability of the transitions. Thus, we asked if we could identify the ideal team structure in a given network, from which we might be able to infer the emergent dynamics. Previously, we had used clustering of the influence matrix to coerce a given network into a two-team configuration. However, we found such clustering to be imperfect for identifying teams. Thus, we sought to devise a different method to estimate teams borrowing from the theory of structural balance. Structural balance theory was first developed by Heider [18, 19], with the hypothesis that psychological systems tend to organize themselves towards lower conflict and higher stability. Thus, the corresponding interaction networks can be such that no triad (three connected nodes) can have an odd number of negative interactions. This theory was further developed by Cartwright and Harary [20], and one of their discoveries was that structurally balanced networks can be partitioned into exactly two “cliques” or groups of nodes, such that all interactions within a group are positive and all interactions across are negative. Thus, networks with two mutually inhibiting nodes are structurally balanced. Further work in structural balance theory identifies that the signs of the elements of the principal eigenvector of the adjacency matrix (corresponding to the eigenvalue with the largest positive real component) follow the groups, such that nodes in the same group have the same sign. Although these results are primarily derived for undirected, signed graphs, we hypothesized that the spectral classification could apply to the case of signed directed networks, as analyzed in the current manuscript. In fact, we find that in two-team networks with mutual inhibition, teams can be identified based on the principal eigenvector - coefficients of the nodes belonging to the same team have the same sign (**Fig 6A, left**). Interestingly, we find that this result can be extended to three team networks by using two eigenvectors instead of one (**Fig 6A, right**). The first eigenvector partitioned one team from the remaining two, which were further partitioned by the second eigenvector. We then extended this analysis to networks with a larger number of teams and asked how well the top eigenvector differentiates the teams. We generated multiple networks of *n* ∈ {2, 3, 4, 5, 6} teams with five nodes per team for varying densities, and calculated the top eigenvectors in each case. We then calculated the average fraction of teams recovered (see methods) by the positive and negative coefficients separately, and found that 50% of the teams were recovered by either segment of the eigenvector (**Fig 6B**). While at low density, the nodes from the same team showed opposite signs in the top eigenvector, the fraction of such broken teams reduced with increasing density (**Fig S7A**). Given these results, we were encouraged to leverage the behavior of principal eigenvector(s) to predict the number of teams. We describe the algorithm in detail in Methods. Briefly, we recursively calculate the principal eigenvector of the adjacency matrix, using the signs of the elements to partition the network into two teams. For the partition that does not follow the definition of teams, where all the edges within the team are positive, we further calculate the principal eigenvector. We created 100 each of *n* ∈ {2, 3, 4, 5, 6} team networks of team size 5 with varying densities, and predicted the teams using this algorithm. For simplicity, we maintained symmetry in team sizes of the network. Despite the directed nature of these networks and the significant disruption of structural balance in networks with a larger number of teams, we found reasonable success in predicting teams. Team prediction was the most accurate in two-team networks, as a direct consequence of their structural balance, with two teams being recovered even at low densities. Density had minimal effect on the recovering teams from two team networks, at different team sizes as well (**Fig S7**). For 3 teams onwards, we find the algorithm predicting a larger number of teams (**Fig 6C**), owing to the fraction of broken teams in the top eigenvector for *n >* 2. Consequently, the ratio of predicted teams to actual teams reduces with increasing density, and increases with increasing number of teams. In three-team networks, two out of the three teams (0.66) are recovered on average at all densities (**Fig 6D**). However, this ratio is much lower for a larger number of teams. Interestingly, this behavior does not depend on the number of nodes, but only on the team configuration, as seen by applying the algorithm to networks with 10 nodes per team (**Fig S7B, C**). In fact, the accuracy improved with increasing number of nodes in the network (compare **Fig 6B, C** with **Fig S7B, C and D, E**). Despite the limited accuracy in identifying the exact teams, we find that the fraction of predicted teams being false positive (i.e., nodes of the predicted team belong to more than one expected teams) is very low (**Fig 6D**), and falls to zero at larger network sizes (**Fig S7F, G**). The teams’ networks become more symmetric as the density of the network increases. We then asked if increasing symmetry would improve the team’s prediction, but we have been unsuccessful. One way we tried increasing symmetry was to use the signed influence matrix [8, 10] instead of the adjacency matrix. Briefly, the influence matrix is a weighted sum of matrices *Adj*^*l*^ and thus has higher density than the adjacency matrix (**Fig S8F**). However, we found that the prediction accuracy was poor for a higher number of teams, with detection of many false positive teams (**Fig S8A-E**). We also attempted to predict teams with the Gramian matrix (*Adj Adj*^*T*^), which is a symmetric matrix. Similar to the case with the influence matrix, the top eigenvector could identify the team separation in two and three-team networks (**Fig S9A-B**), the prediction accuracy was poor for more than three teams (**Fig S9C-E**). Interestingly, however, for three teams, the Gramian matrix predicted teams better than the influence matrix. We further evaluated the team configuration in the influence and Gramian matrices for different densities and found that, especially in networks with more than three teams, the inconsistency with team configuration was high, which could explain the poor prediction accuracy of these two matrices (**Fig S8G, S9G**). Therefore, we conclude that a successful classification would require a transformation of the adjacency matrix that maintains the team configuration and has high density and symmetry. Our results suggest that it is possible to extend the spectral properties of structurally balanced networks to directed, multi-team networks as well, prompting further analytical work.

**Figure 6.**
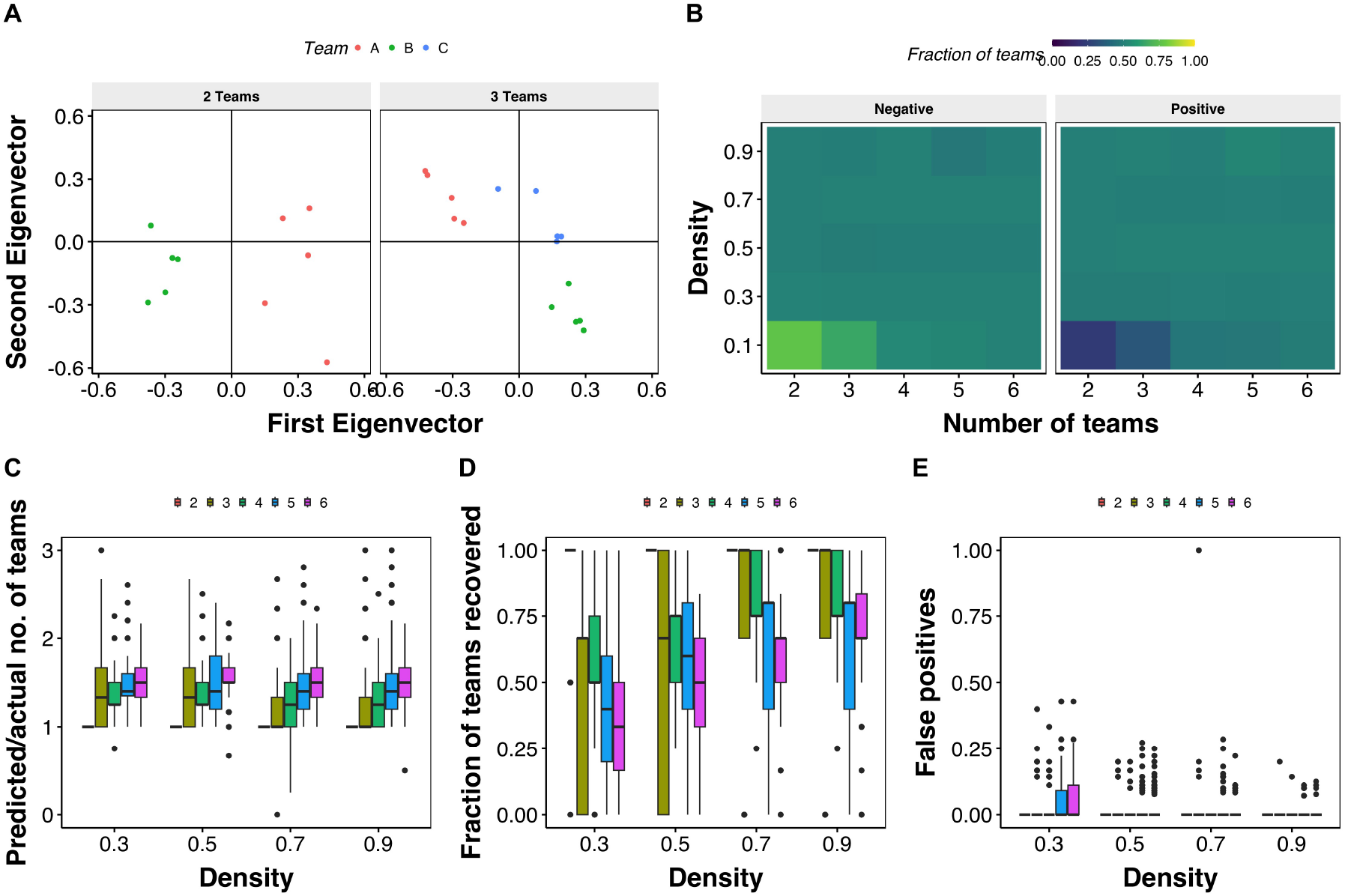
Spectral properties can partition a multi-team network. **A** Scatterplot depicting the weights of each node in the eigenvectors corresponding to the highest (principal eigen vector) and second-highest positive eigenvalue. Each point corresponds to a node, and the team identity is labelled by the color and shape of the point. Note that all nodes belonging to the same team have the same sign in the principal eigenvector, a property we used to identify teams. **B** Heatmaps depicting the mean fraction of teams recovered in the top eigenvector with positive coefficients and negative coefficients across 100 iterations each for different combinations of density and number of teams. **C** Boxplot depicting the change in the ratio of the number of teams predicted by the algorithm to expected teams, as a function of density (x-axis) and the number of teams (color). **D** Same as B, but for a change in the fraction of teams recovered. **E** Fraction of false positive predictions, where a false positive is identified as a predicted team having nodes from more than one expected team

**Figure 7.**
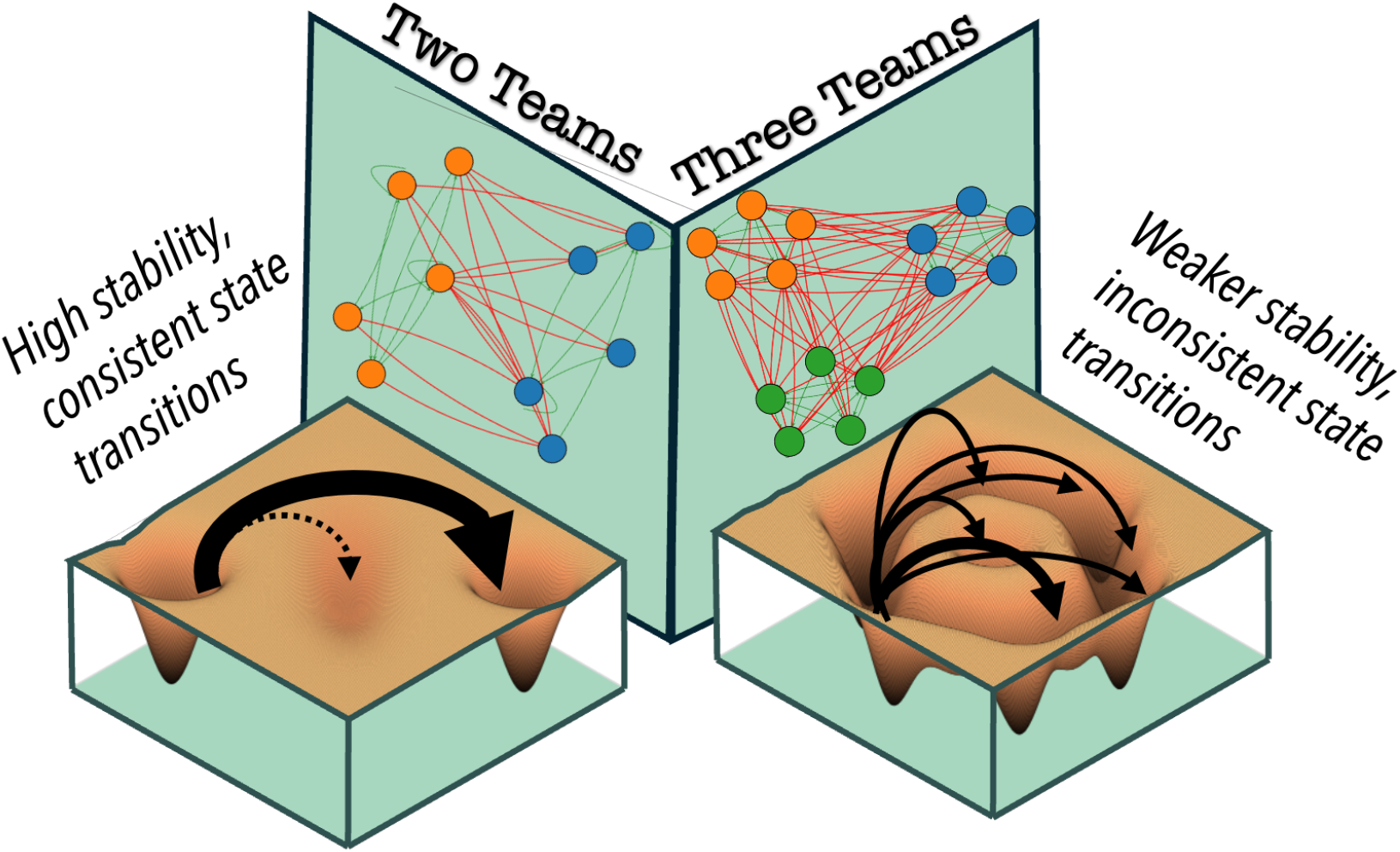
Schematic demonstrating the inefficiency of multi-team systems as decision systems.

## 3. Discussion

Our study addresses a fundamental question in biology: why are binary cell-fate decision-making systems far more prevalent than multi-fate systems? By extending the well-characterized gene regulatory networks (GRNs) underlying binary decisions—composed of two mutually inhibitory “teams” of nodes — to networks with three or more such teams, we uncovered the functional advantages that binary systems might have over multi-fate systems. Our findings robustly demonstrate that multi-team networks are inherently less stable and unreliable for cell-fate decision-making compared to two-team (binary) networks, providing a compelling explanation for their scarcity in biological systems. A central finding is the significant reduction in the stability of “pure” (single positive) states in three-team networks, with their complete lack in four teams and beyond. Single positive states studied here represent functionally distinct phenotypes that allow biological systems to be modular. Therefore, attaining pure states is a highly desirable property of any cell-fate decision system. While three-team networks are also capable of giving rise to single-positive steady states, their stability is lower, and they can only be achieved at higher densities [17]. This reduction in stability can be attributed to an increased “frustration” in the network, which can be measured by an increase in the fraction of negative feedback loops with an increasing number of teams. We demonstrate that the steady states emergent from multi-team networks are more frustrated than those from two-team networks, measured as the fraction of edges in the network whose signs are not satisfied by the configuration of the participating nodes. This increased frustration also leads to a decrease in the stability of the steady states.

The stability of two-team networks demonstrated here provides insights into biological processes like dedifferentiation and transdifferentiation. The robustness of two-team networks makes the corresponding binary cell-fate decision systems the perfect building blocks of progressive differentiation. It follows that dedifferentiation (reprogramming), which requires reversing multiple such binary decisions, demands precise, large-scale perturbations across several two-team networks, explaining its low success rate [21]. In contrast, transdifferentiation, which involves switching between two phenotypic states, typically requires the control of most two-team networks, making it relatively easier to achieve [22]. However, the discovery of Yamanaka factors, which enable reprogramming to pluripotency, suggests additional network nuances, such as feedback loops or asymmetric interactions, that warrant further investigation [23]. These factors highlight the limitations of our Boolean formalism, which, while computationally efficient, simplifies the continuous dynamics of real biological systems. While single positive phenotypes are important, multi-positive or hybrid phenotypes also have significance in various contexts. Stem cells have been shown to have higher entropy as compared to differentiated cell types [24, 25], indicating that stem cells are more hybrid in their expression patterns. Other instances of hybrid phenotypes include hybrid E/M phenotypes in the context of cancer metastasis, which express a mixture of genes that identify the terminally differentiated Epithelial and Mesenchymal phenotypes [26]. Hybrid E/M phenotypes have been associated with improved metastatic potential due to enhanced stemness, immune evasion, and drug resistance [27]. In both cases, prioritizing the stability of these hybrid phenotypes is not desirable, since high stemness is not conducive to normal functionality of organisms, and high hybridness would increase the incidence of metastasis. Thus, it is more likely for two-team networks to control the decision-making involving such hybrid phenotypes rather than multi-team networks. We previously demonstrated that increasing the fraction of negative feedback loops in a network can give rise to a higher frequency of hybrid states [28]. A generalization of this principle can be achieved by introducing “impurities” in two-team networks (flipping the sign of a fraction of edges in ideal two-team networks), which has been shown to increase the frequency of hybrid phenotypes, especially at a density of 0.2-0.3, where many of the real-world GRNs lie [29]. Such impurities are naturally found in networks across contexts and disciplines. A future direction for the current manuscript would be to explore the effects of impurities in multi-team competition dynamics.

Coexistence of nodes from multiple teams is significant in other contexts, such as social and ecological networks. In ecological networks where a fundamental competition for resources exists for all species sharing an ecosystem, the coexistence of multiple species is often seen. Our model suggests that such coexistence can emerge from multi-species competition (double positive states in three-team networks, for example), but has limited stability under a noisy environment. The stability must therefore be a product of various other factors, such as the development of cooperative interactions, spatial isolation, and resource heterogeneity [30, 31]. Thus, another extension of our current work can be to construct multilayer networks that incorporate the different aspects of social interactions and their effect on steady states and transitions. Interestingly, a recent analytical study on the competition-colonization trade-off predicts the average number of species surviving at any given time to be roughly half of the total number of species, a result similar to our analysis of four and five-team networks [32].

Our results align with Heider’s theory of structural balance, which posits that networks evolve toward configurations where the number of negative interactions (inhibitions) in any triad is even (called balanced triads), minimizing conflict [14, 18]. Two-team networks inherently satisfy this property as all within-team interactions are positive, and between-team interactions are negative, leading to all the triads being balanced. In contrast, three-team networks exhibit intrinsic frustration due to unavoidable negative triads, correlating with their reduced stability and heightened perturbation sensitivity. This structural balance can explain the prevalence of binary competition in both biological and social systems, as networks may evolve toward balanced configurations to maximize stability. Multi-team competition extends far beyond biological systems, manifesting across diverse domains where multiple entities compete for resources, influence, or dominance. In political systems, multi-party democracies often experience instability and coalition fragmentation, frequently consolidating into effective two-party systems despite initial multi-faction landscapes [13]. Similarly, in economic markets, oligopolistic competition frequently evolves toward duopolistic structures, suggesting inherent stability advantages in binary competitive frameworks. Social media platforms exhibit analogous patterns, where complex multi-group discussions often polarize into binary opposing camps, despite initial diversity of viewpoints [33, 34].

Notably, our findings are for directed signed networks, extending the applicability of structural balance beyond undirected graphs. Spectral analysis further supports this, as the principal eigenvector accurately partitions two-team networks, with reasonable success for three-team networks (and beyond). We would like to highlight that the spectral classification presented here is different from the standard spectral clustering methods used to identify communities in networks. The primary difference is, the methods of spectral clustering identify communities that are primarily defined for either signed or directed graphs [35, 36]. Analytical understanding of the spectral properties of directed signed graphs is heavily lacking, due to various factors, including the asymmetry of the adjacency matrix. The fact that GRNs have an abundance of self-loops further complicates their spectral properties. Some recent studies proposed alternatives to the commonly used Laplacian matrix, such as magnetic Laplacians that transform the adjacency matrices of DSG to achieve a symmetric matrix [37]. But we find that team structure does not require such transformations for meaningful community detection. Another primary difference between the standard spectral clustering problem and team classification is that the communities identified in spectral clustering are well connected within themselves but are sparsely connected across, thereby leading to “modular” communities. Here, however, the connectivity can (and does in this scenario) remain the same between teams, and in some cases exceeds the connectivity within a team, such as teams formed from the logic of “enemy of enemy is a friend” (see networks in [10].

Feedback loops play crucial roles in determining the emergent behavior of networks. Positive feedback loops can lead to signal amplification, hysteresis and corresponding memory, ultrasensitivity, and multistability [38–40], while negative feedback loops can lead to a wide range of behaviors from chaos, to different kinds of oscillations to robust mono-stability (e.g., perfect adaptation in bacterial systems) [41–43]. Complex behaviors can emerge when positive and negative feedback loops interact with each other [44–48]. A relevant example is a toggle triad. While a repressilator is a perfect model of an oscillating system, a toggle triad gives rise to single-point attractors [49]. In GRNs with multiple interacting negative and positive feedbacks, negative feedback has generally been associated with weaker stability and frustration in the state space, while positive feedback has been associated with (multi)stability [50]. The presence of negative feedback loops causes the unbalanced triads to break structural balance. Therefore, two team networks without any impurities do not have any negative feedback loops. Three teams and beyond, on the other hand, are bound to have negative feedback loops. Any feedback involving nodes from an odd number of teams (e.g., 4 4-node feedback loop with two nodes in one team and the third and fourth node in a different team each) will be a negative feedback. Interestingly, while the number of such negative feedback loops increases as the density of multi-team networks increases, the state space becomes more stable at higher density. While it is tempting to explain this behavior away by highlighting that the positive feedback in the network increases as well, and somehow overpowers the instability created by the negative feedback, the ratio of positive to negative feedback loops should not change significantly in lower densities as well, invalidating this explanation. Thus, while the increasing negative feedback from two to three teams and beyond does have a role to play in decreasing stability, the multi-team networks provide a unique opportunity to investigate how interactions between positive and negative feedback give rise to complex stability features.

Our study has certain limitations. The use of a Boolean formalism, while computationally efficient for simulating large networks, is a simplified representation of the continuous dynamics of real biological systems. While our results are supported by investigations of smaller systems using continuous simulation formalisms [49], and the symmetry in our networks, the effect of different simulation formalisms on our results has to be quantified. Our networks were artificially generated based on a specific “mutually inhibitory teams” motif, which may not encompass all possible architectures for multi-fate biological systems. Future research could explore whether different underlying network architectures, beyond the direct extension of this motif, might inherently support more stable multifate decisions. In the context of social and ecological networks, studying multi-team competition in the presence of external factors such as spatial isolation or dynamic interactions can provide further insights into understanding how stability can be achieved in these contexts. Further validation on experimentally derived GRNs of actual multi-fate systems, such as the T-cell differentiation network mentioned, would strengthen the direct applicability of these theoretical findings to real biological processes.

In conclusion, our findings strongly suggest that the prevalence of binary cell-fate decisions in biology is not accidental. The inherent instability, high frustration, unpredictable transitions, and unreliability under targeted perturbations of multi-team gene regulatory networks make them evolutionarily less favorable. For processes critical to development, disease (e.g., preventing aberrant cell fates in cancer or metastasis), and regeneration, the robustness and predictability offered by two-team networks provide a significant functional advantage.

## 4. Methods

### 4.1. Notation

The following notations are followed throughout the article unless mentioned otherwise:

- *N* : Number of nodes in a network
- *E*: Number of edges in a network
- Adj: Adjacency matrix
- *i, j*: Indices that refer to the nodes in the corresponding positions in adjacency matrices
- Adj_*ij*_ : The interaction strength of the edge from the *i*th node to the *j*th node

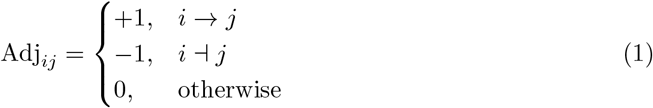
- *S*(*t*): State of a network at time *t*
- *s*_*i*_(*t*) ∈ {−1, 1}: State/activity of a node of a network at time *t*

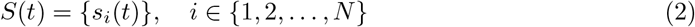

### 4.2. Figure 1

#### 4.2.1. Boolean Simulations

Boolean modeling is a parameter-independent way of modeling biological networks with lesser computational costs compared to ODE formalisms, the reduced computational costs enables us to stimulate and study larger networks. We use a threshold based Boolean formalism in which each node of the network is a binary variable (−1 or 1) and the interactions between the nodes are modeled as edges with two values ((−1 or 1) depending on the nature of interactions (excitatory or inhibitory). A state of a network is defined by a binary string of variables *s*_*i*_, which gives information about which node is active/ON (*s*_*i*_ = 1) or inactive/OFF (*s*_*i*_ = −1). The interactions between the nodes(edges) are represented and stored using the nonsymmetric adjacency matrix Adj, where each element of the matrix, Adj_*ij*_, is the interaction strength of the edge from the *i*th to *j*th node of the network. All activations are given a weight of +1, and all inhibitions are given weight of −1.

The simulations are conducted asynchronously (one randomly chosen node is updated at each iteration). The state of the system is updated using a majority rule given below:

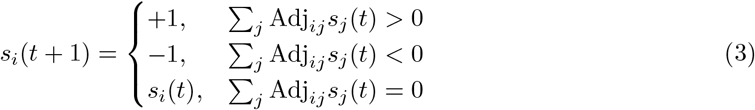

Simulations are carried out until either of the two conditions is reached:

1. *t >* 1000
2. *s*_*i*_(*t* + 1) = *s*_*i*_(*t*) for all *i* ∈ {1, …, *N* }

The latter condition implies that a steady state has been reached. *S*(*t*) is identified as the steady state.

#### 4.2.2. Network Generation

We generated artificial networks with teams using Stochastic block model [**SBM**]. For a network of *n* teams, with each team having *m*_*i*_, *i* ∈ 1, 2, …, *n* and a uniform density *d*, we start by creating an empty adjacency matrix of the dimensions Σ*m*_*i*_ *X* Σ*m*_*i*_. This matrix is then divided into *n*^2^ submatrices/blocks with the dimensions *m*_*i*_ *X m*_*j*_ : *i, j* ∈ 1, 2, …, *n*. We then randomly select [*m*_*i*_ * *m*_*j*_ * *d*] cells in each block, set their values to 1 if *i* = *j* and −1 if *i* ≠ *j*. The value of the remaining *m*_*i*_ * *m*_*j*_ − [*m*_*i*_ * *m*_*j*_ * *d*] cells is set to 0.

#### 4.2.3. Fraction of Single Positive States

Each network generated is simulated using the Boolean formalism starting from 10,000 initial conditions and the steady states are stored, out of these steady states, the fraction of states which have only one Team in On position (+1) and all other teams Off (-1) count towards the fraction of single positive steady states

### 4.3. Figure 2

#### 4.3.1. Frustration

Frustration is a measure of the agreement between the network topology and a given steady state. For a given network and state, the frustration is calculated as follows:

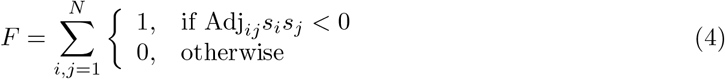

where *E* is the number of edges in the network.

#### 4.3.2. Number of Teams which are On

Each network generated is simulated using the Boolean formalism starting from 10,000 initial conditions, and the steady states are stored, out of these steady states, the fraction of states which have any combination of n Teams in On position (+1) and all other teams Off (-1) count towards the fraction of states with N Teams On.

### 4.4. Figure 3

#### 4.4.1. State classification

For each network in our ensemble, we classify non-hybrid initial and destination states into the types ‘+’ (10,01 in two team networks and 100,010,001 in three team networks) and ‘++’ (110,101,011 in three team networks only). On top of this scheme, destination states are further sub-classified as ‘mirror’ if they have a hamming distance of 1 from their corresponding initial states or ‘non-mirror’ otherwise.

#### 4.4.2. Untargeted simultaneous perturbation

We start with a steady state of the network *S*_0_. Let N be the set of nodes in the network. We fix the number of nodes we intend to perturb as k, and construct a set V by randomly sampling k nodes from N without replacement. We construct a new state *S*_1_ such that for each node i in the network,

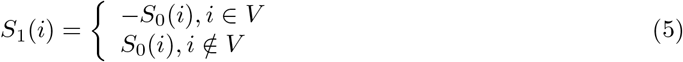

We evolve *S*_1_ using the evolution algorithm described in Eq. 3 to get steady state 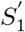.

For each initial state classified as ‘+’ or ‘++’ on a given network, we perform the experiment with levels of perturbation ranging from a single node to all nodes in the network (with increments of 1 node). For each specific initial condition and level of perturbation on the network, 100 repetitions were carried out. For each repetition we record the destination after switching. We proceed to find the frequency of each class (+ mirror/+ non-mirror/ ++ mirror /++ non-mirror) of destinations over all repetitions for each class (+/++) of initial conditions and each level of perturbation on the network.

#### 4.4.3. Untargeted sequential perturbation

We start with a steady state of the network *S*_0_. Let N be the set of nodes in the network and let *V*_*t*_ be the set of perturbed nodes at time t, with *V*_0_=.At time *t*, we pick a single node j at random from the set N *V*_*t*−1_ and construct *V*_*t*_=*V*_*t*−1_ ⋃ {j }. We construct a new state *S*_1_ such that for each node i in the network,

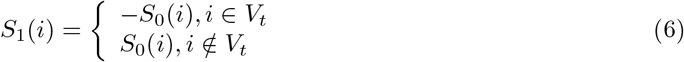

We evolve *S*_1_ using the evolution algorithm described in Eq. 3 to get steady state 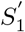. We proceed to the next time-step *t* + 1 if 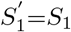; else we end the experiment at time *t*.

For each initial state classified as ‘+’ or ‘++’ that is stable on a given network, we perform 100 repetitions of the experiment. For each repetition we recorded the total perturbation needed for switching and the destination after switching. We average the total perturbation needed and the hamming distance between initial and final states over all repetitions for each class of initial conditions on the network. We also find the frequency of each class of destinations over all repetitions for each class of initial conditions on the network.

### 4.5. Figure 4

#### 4.5.1. Directed simultaneous perturbation

Let a network have *n* teams (known a-priori). Let *N*_*k*_ be the set of nodes in the *k*^*th*^ team and *c*_*k*_ be the number of nodes to be perturbed in the *k*^*th*^ team. For each team *k*, we create set *V*_*k*_ by randomly sampling *c*_*k*_ nodes from *N*_*k*_ without replacement. We construct a new state *S*_1_ such that for each node i in the network,

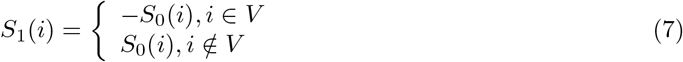

where 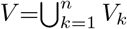

We evolve *S*_1_ using the evolution algorithm described in Eq. 3 to get steady state 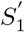.

For each network in our ensemble, we performed the directed simultaneous perturbation experiment with all combinations of directed perturbation to the initial condition 10 on two-team networks and the conditions 100 and 110 on three-team networks. A directed perturbation for a network of n teams is 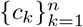 with *c*_*k*_ ∈

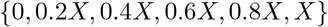

where X is the number of nodes in the network and each *c*_*k*_ is independent. For each combination of initial condition and directed perturbation on the network, 100 repetitions were carried out. For each repetition we record the destination after switching. We proceed to find the frequency of each destination over all repetitions for a given combination of initial condition and directed perturbation on the network.

#### 4.5.2. Directed sequential perturbation

The procedure is identical to that of undirected sequential perturbation, with the only difference being that *N* can now be any subset of nodes in the network.

A destination direction is the steady state which we wish to perturb the system in the direction of. For a given initial state *S*_0_ and destination direction *S*_1_, we construct *N* as the set of all nodes *i* such that *S*_1_(*i*) ≠ *S*_0_(*i*) ∀*i* ∈ *N* . For each network in our ensemble, we performed the directed sequential perturbation experiment with the following initial conditions and destination directions:

1. Two-team networks:
  a. **Initial**: 10, hybrid
  b. **Direction**: 01
2. Three-team networks:
  a. **Initial**: 100, 010, 110, 101, hybrid
  b. **Direction**: 001, 011

For each combination of initial condition and destination direction on a network, 100 repetitions of the experiment were performed. Hybrid initial conditions were separately generated for each network by generating a single random initial condition per network which was neither single/double positive and which was stable on the given network. All 100 repetitions on the same network for ‘hybrid’ used the same initial condition. For the experiment, only nodes whose states differed between the initial condition and destination direction were allowed to be flipped. For each repetition we recorded the total perturbation needed for switching and the destination after switching.

We fit the resultant data to a Cox Proportional Hazards Model. Each repetition constitutes an individual data entry characterised by 6 Boolean variables which can take values of 0 (no) or 1 (yes) depending on whether the entry satisfies the corresponding criteria - ‘# teams = 2’, ‘# teams = 3’, ‘initial state = +’, ‘initial state = ++’, ‘initial state = mirror (relative to destination direction)’, ‘initial state = hybrid’. These 6 variables constitute our input data. For our output data, we compute the variables ‘number of flips required for switching’ and ‘destination = destination direction’ (the latter variable is Boolean) corresponding to each input data entry. We train the model on this data. This was performed on python using the CoxPHSurvivalAnalysis() function from the scikit-survival (sk-surv) package.

### 4.6. Figure 6

#### 4.6.1. Framework for scoring; notation

We tested our algorithm on randomly generated networks. A randomly generated network on *N* nodes is a function of the number of teams (*N*_*t*_), the size of each team (*T*_*s*_), and the density (*d*_*s*_), where *N*_*t*_ and *d*_*s*_ are constants, and *T*_*s*_ is a list of size 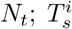 corresponds to the size of the *i*-th team. The density of edges within a team and edges leaving any team is taken to be the same, fixed to *d*_*s*_. The number of edges within and leaving team *i* depends on 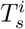 and *d*_*s*_; and these edges are distributed randomly.

The algorithm outputs predictions as a set of lists. The number of lists within the set are the predicted number of teams (*PN*_*t*_) in the input network. Each of these lists contains the node labels that belong to the same team; the *i*-th list in the prediction set is the *i*-th predicted team 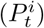. Note that *PN*_*t*_ need not be the same as *N*_*t*_. To assess the performance of the model, we will use the metrics described in the next subsections.

#### 4.6.2. Algorithm

#### 4.6.3. Calculation of the influence matrix

The influence matrix is a transformation of the adjacency matrix that attempts to incorporate indirect interactions between the nodes of a network, and is calculated as follows:

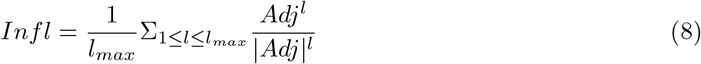

where *Adj* is the adjacency matrix, |*Adj*| represents the magnitude of the *Adj*, and *l*_*max*_ is the maximum path length considered, typically 10.

#### 4.6.4. Metrics to measure the success of team prediction

##### Predicted / actual number of teams

The ratio of the number of teams predicted to the ground truth number of teams

##### Fraction of teams recovered

Among the predicted teams (where each team is a set of nodes), the number of teams that are not proper subsets of the ground truth teams.

##### False positives

The number of predicted teams whose elements belong to different ground truth teams.

###### Algorithm 1

Iteratively Perform Itersort

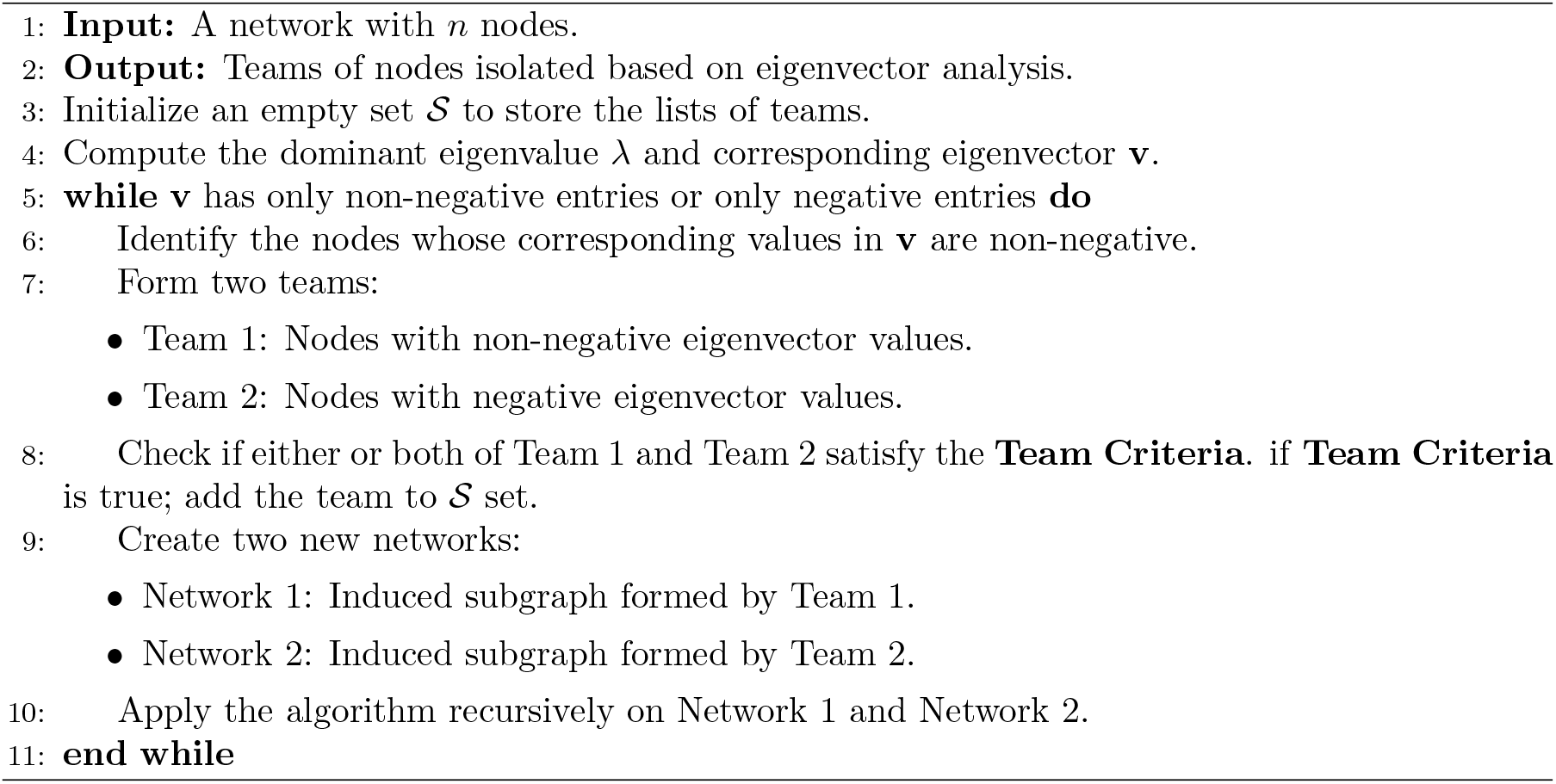

## 5. Funding and Acknowledgements

MKJ was supported by the Ramanujan Fellowship (SB/S2/RJN-049/2018) awarded by the Science and Engineering Research Board (SERB), Department of Science and Technology, Government of India. MKJ was also supported by Param Hansa Philanthropies. KuH, VA and AM were supported by KVPY. KH was partially supported by Prime Ministers’ Research Fellowship (PMRF), Government of India, the National Science Foundation through the Center for Theoretical Biological Physics, PHY-2019745 and under Award Number MCB-2114191. TG was partially supported by NSF grant DMS-1951510. We would like to acknowledge Divyoj Singh and Anish Hebbar for the valuable discussions on the spectral properties of teamed networks.

## 6. Author contributions

Conceptualization : KiH and MKJ. Methodology : KiH, VA, KuH, AM. Software, Formal analysis, Visualization : VA, KuH, AM. Writing and review : all authors

## 7. Data and code availability

All codes used in the manuscript are available at https://github.com/askhari139/nTeamsProject.

