## Supplementary figures and note for "Multi-team conflict resolution is ineffective for stable decision making"

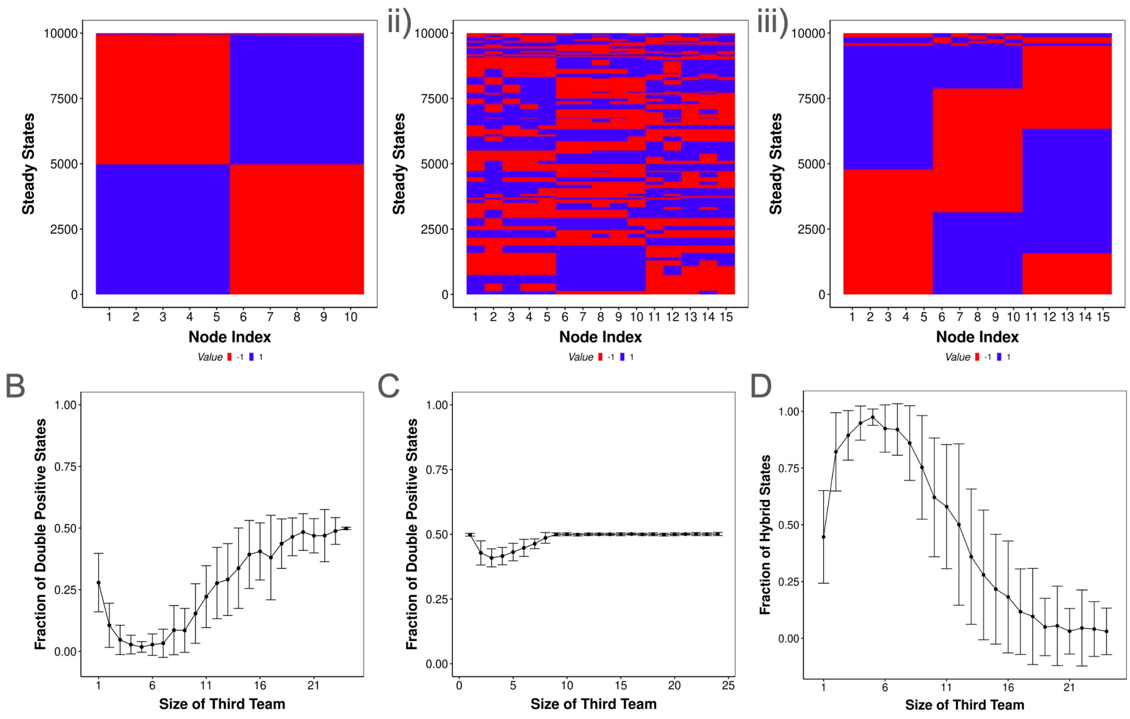


Figure S1: **Steady state frequency distribution of two vs three teams at varying density and team size.** **A** Representative heatmaps depicting the configuration of steady states in (i) two team network at 0.9 density, (ii) three team network at 0.3 density and (iii) 0.9 density. **B** Fraction of double positive states (all nodes from two teams are on, all nodes from the third team are off) with two teams of 5 nodes each and changing size of third team, at an edge density of 0.3. Error bars represent standard deviation over 50 networks. **C** Same as B, but for an edge density of 0.9. **D** Same as B, but for hybrid states expressing nodes that form strict subsets of one or more teams.

### 8. Supplementary Figures

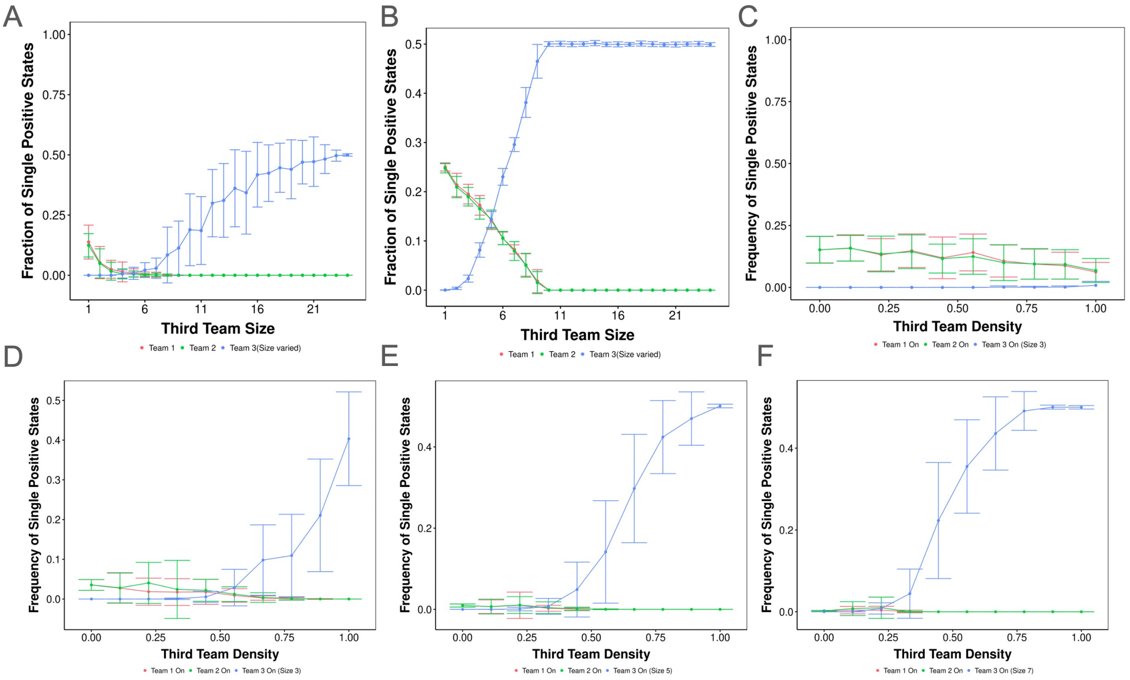


Figure S2: **Specific configurations of single positive states emergent from three team networks**. **A-B** Fraction of single positive states corresponding to each team as the size of the third team is varied for 0.3 and 0.9 edge density. **C-E** Same as A,B but for varying density at for a team three of size 1,3,5 and 7 respectively. In each plot, the color represents the team for which single positive state frequency is reported. Error bars represent the standard deviation obtained over 50 networks.

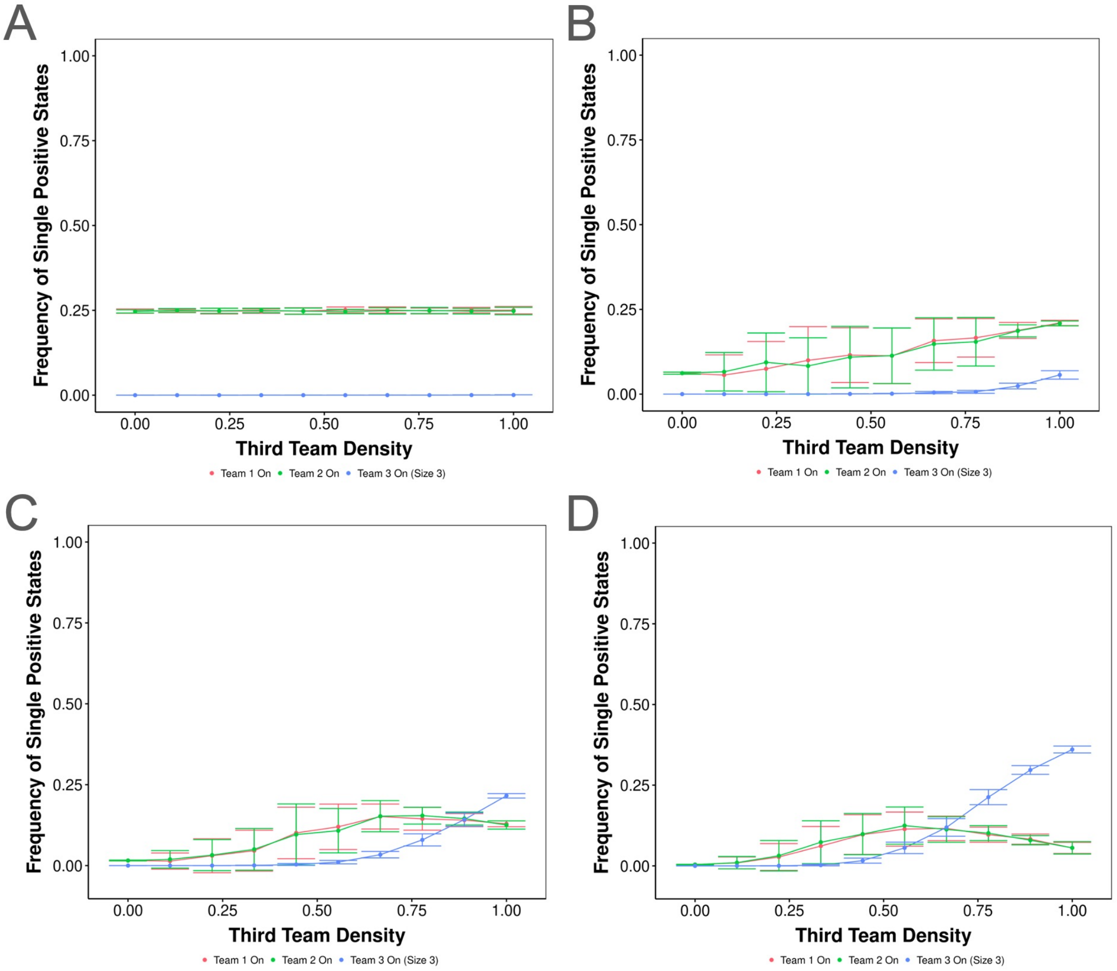


Figure S3: Specific configurations of single positive states emergent from three team networks

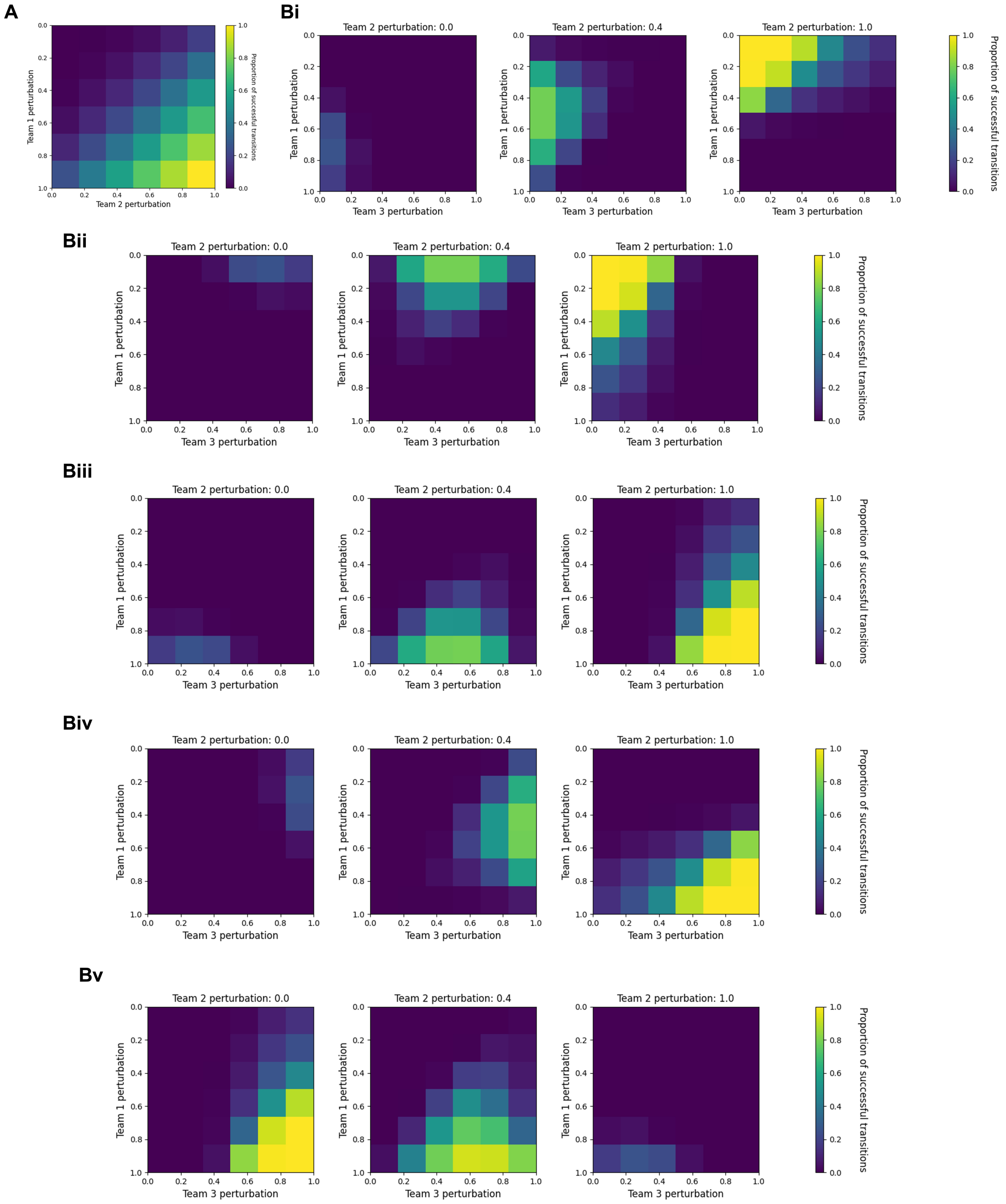


Figure S4: The probability of successful state transitions for directed perturbations in two vs three team networks. **A.** Same as Fig. 4A, but for 0.3 density. **B** Same as Fig 4B, but for

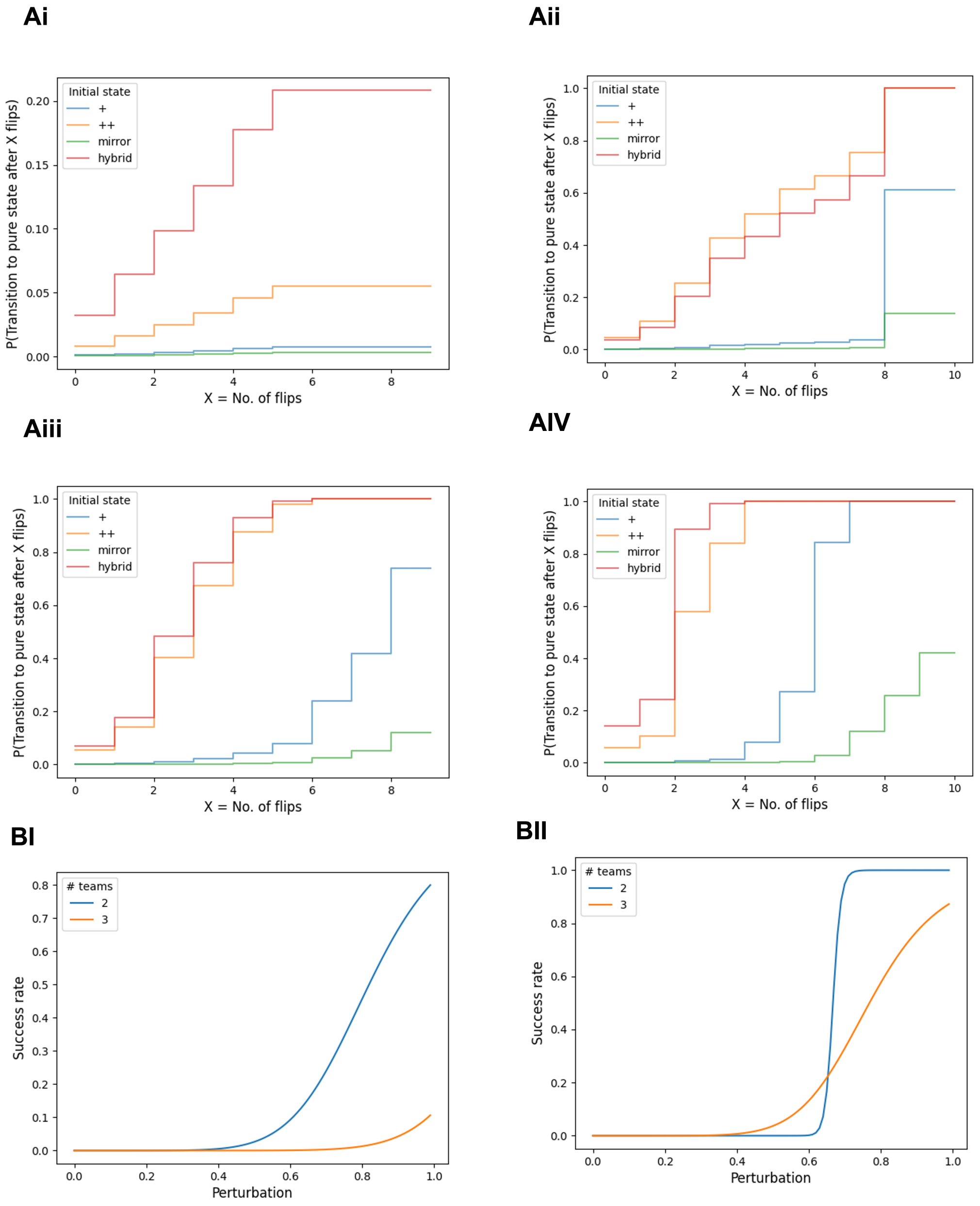


Figure S5: **A**Survival plots depicting the probability of successful transition to a single positive (pure) state, starting from different kinds of initial states in three team networks for increasing density values. **B** Comparison of rate of successful transition from any origin to single positive state upon directed perturbation for (i) 0.3 and (ii) 0.9 density

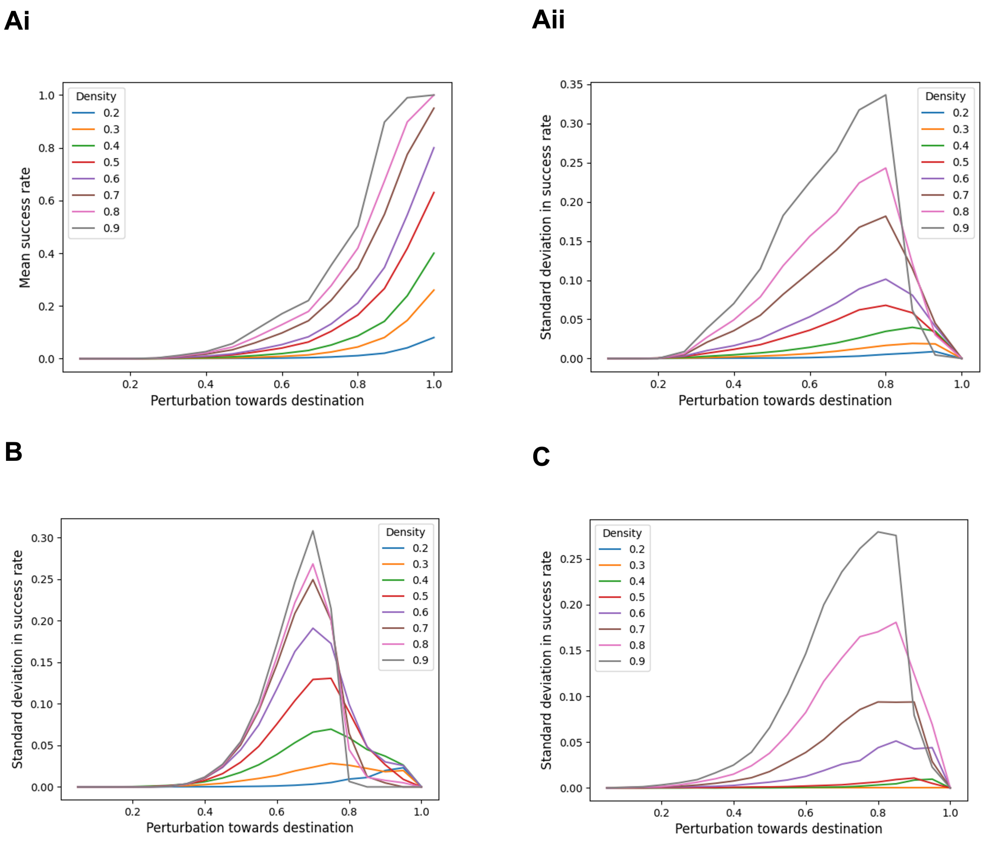
Ai
Density
0.2
0.3
0.4
0.5
0.6
0.7
0.8
0.9
Mean success rate
Perturbation towards destination
Aii
Standard deviation in success rate
Perturbation towards destination
B
Standard deviation in success rate
Perturbation towards destination
C
Standard deviation in success rate
Perturbation towards destination

Figure S6:

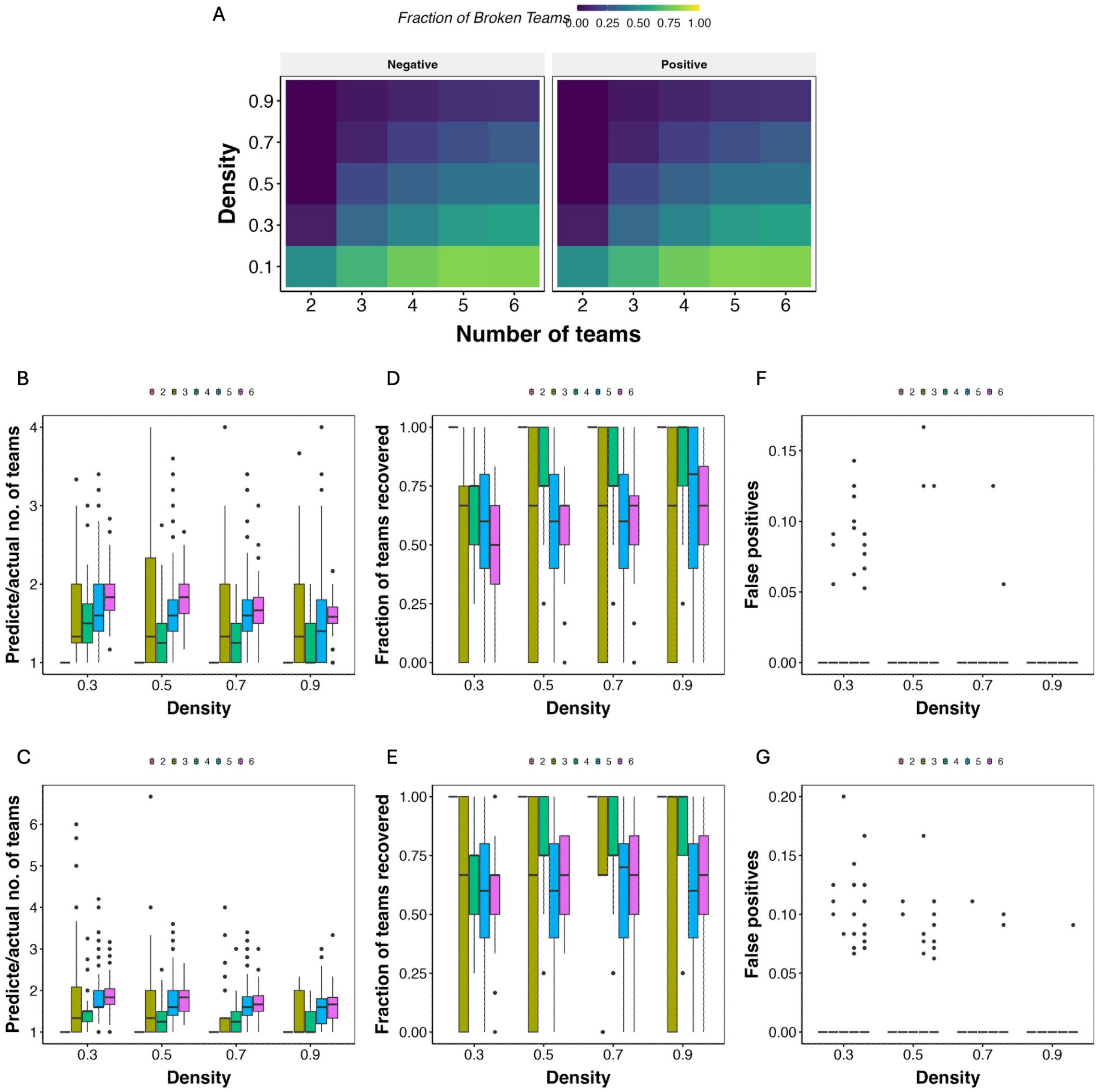


Figure S7: **A** Mean fraction of teams with partially positive/ partially negative coefficients in the top eigenvector of the adjacency matrix for different densities and number of teams. **B-D** Same as **Fig 6 C-E**, but for a teamsize of 10 nodes. **E-G** Same as **B-D** but for teamsize of 20 nodes.

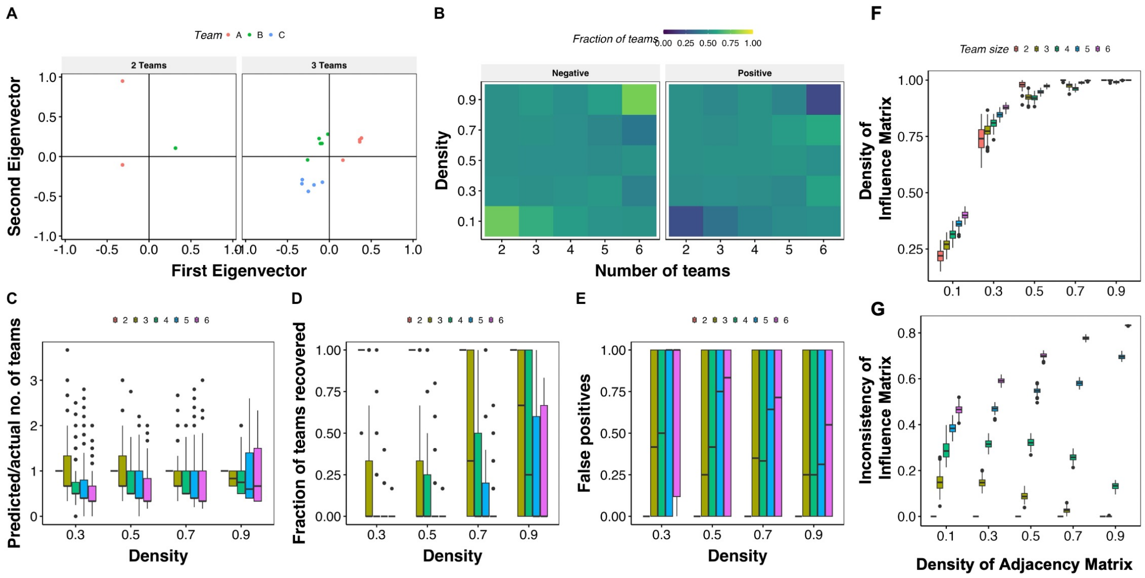

Figure S8: **Team classification using the influence matrix A** Scatterplot depicting the weights of each node in the eigenvectors corresponding to the highest and second highest positive eigen value. Each point corresponds to a node, and the team identity is labeled by the color and shape of the point. **B** Heatmaps depicting the mean fraction of teams recovered in the top eigenvector with positive coefficients and negative coefficients across 100 iterations each for different combinations of density and number of teams. **C** Boxplot depicting the change in the ratio of the number of teams predicted by the algorithm to expected teams, as a function of density (x-axis) and the number of teams (color). **D** Same as B, but for change in the fraction of teams recovered. **E** Fraction of false positive predictions, where a false positive is identified as a predicted team having nodes from more than one expected teams

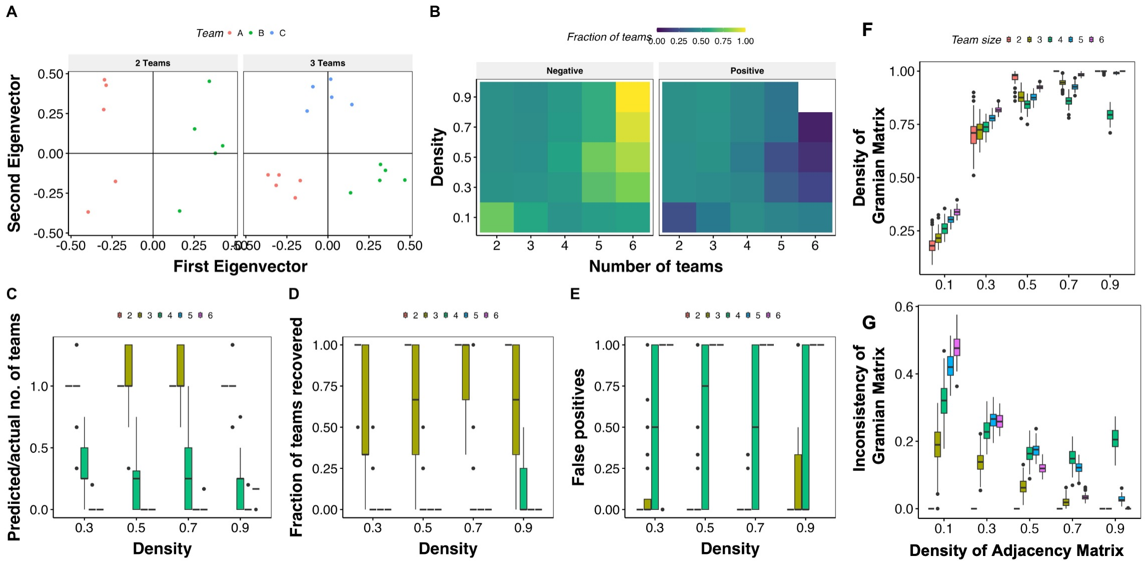


Figure S9: **Team classification using the Gramian matrix A** Scatterplot depicting the weights of each node in the eigenvectors corresponding to the highest and second highest positive eigen value. Each point corresponds to a node, and the team identity is labelled by the color and shape of the point. **B** Heatmaps depicting the mean fraction of teams recovered in the top eigenvector with positive coefficients and negative coefficients across 100 iterations each for different combinations of density and number of teams. **C** Boxplot depicting the change in the ratio of the number of teams predicted by the algorithm to expected teams, as a function of density (x-axis) and the number of teams (color). **D** Same as B, but for change in the fraction of teams recovered. **E** Fraction of false positive predictions, where a false positive is identified as a predicted team having nodes from more than one expected teams

### 9. Analytical results

#### *9.1. Maximum frustration of steady states in ising formalism*

An edge $ij$ is said to be frustrated for a given state $S(t)$ (not necessarily steady state), if

$$Adj_{ij}s_i(t)s_j(t) < 0 \tag{9}$$

Correspondingly, the frustration of a state is defined as follows:

$$f_S(t) = \frac{\sum_i \sum_j 1_{\mathbb{Z}^-}(Adj_{ij}s_i(t)s_j(t))}{\sum_i \sum_j |Adj_{ij}|}, \quad further$$
$$\sum_i \sum_j |Adj_{ij}| = \sum_i T_{1_i} + T_{2_i} \quad where$$
$$T_{1_i} = \sum_j 1_{\mathbb{Z}^-}(Adj_{ij}s_i(t)s_j(t))$$
$$T_{2_i} = \sum_j 1_{\mathbb{Z}^+}(Adj_{ij}s_i(t)s_j(t))$$

Thus,

$$f_{S(t)} = \frac{\sum_i T_{1_i}}{\sum_i T_{1_i} + T_{2_i}} \tag{10}$$

In the ising boolean formalism, the update rules are defined as follows:

$$s_i(t+1) = \begin{cases} +1, \sum_j Adj_{ij}s_j(t) > 0 \\ -1, \sum_j Adj_{ij}s_j(t) < 0 \\ s_i(t), \sum_j Adj_{ij}s_j(t) = 0 \end{cases} \tag{11}$$

with standard definitions of all involved terms. The update rules indicate that, for the activity of a node to remain conserved in the next time step,

$$s_i(t+1) = \begin{cases} \sum_j Adj_{ij}s_j(t) >= 0, s_i(t) = +1 \\ \sum_j Adj_{ij}s_j(t) <= 0, s_i(t) = -1 \end{cases} \tag{12}$$

In other words,

$$s_i(t+1) = s_i(t) \ if \ \sum_j Adj_{ij}s_j(t)s_i(t) >= 0 \tag{13}$$

Thus, the condition for a state $S(t)$ to be a steady state is:

$$\sum_j Adj_{ij}s_j(t)s_i(t) >= 0 \forall i \tag{14}$$

It is possible to establish a relationship between frustration and the possibility of a state being stable. Now, consider the following two extreme cases of states:

- All edges are frustrated, i.e., $Frustration = 1$. In this case, Since $Adj_{ij}s_i(t)s_j(t) < 0$ for all $i, j$, the condition in 13 is never satisfied. Hence, none of the nodes in the state can retain their activity, and therefore the state cannot be a steady state.

- No edge is frustrated, i.e., $Frustration = 0$. In this case, the condition in 13 is satisfied for all $i, j$ and hence for all nodes, making the state steady.

These extreme cases hence seem to indicate that the higher the frustration, the lesser the chance of a state being steady. However, having the frustration value as 1 is seldom possible, due to the presence of signal nodes (not influenced by any other node in the network and therefore can retain any activity), negative feedback, and feed-forward loops. Thus, there exists a threshold value of frustration above which a state ceases to be a steady state.

Note that the magnitude of the term $Adj_{ij}s_i(t)s_j(t)$ is always 1. Therefore, the condition for the stability of the $S(t)$ (Equation 14) can be re-written as follows:

$$\begin{aligned} T_{1_i} - T_{2_i} &\leq 0 \forall i \\ i.e.,\ T_{1_i} &\leq T_{2_i} \forall i \\ i.e.,\ \sum_i T_{1_i} &\leq \sum_i T_{2_i} \\ \frac{\sum_i T_{2_i}}{\sum_i T_{1_i}} &\geq 1 \end{aligned} \tag{15}$$

Rewriting 10, we get

$$f_{S(t)} = \frac{1}{1 + \frac{\sum_i T_{2_i}}{\sum_i T_{1_i}}} \tag{16}$$

Note that $\sum_i T_{1_i} = 0$ ensures that there are no negative elements in LHS of Equation 14 and thus ensures the stability of $S(t)$. Similarly, this condition sets $f_{S(t)} = 0$. Hence, in the above equations, we can use $\sum_i T_{1_i} \neq 0$ without loss of generality. Using 15, we have

$$\begin{aligned} \frac{1}{f_{S(t)}} &= 1 + \frac{\sum_i T_{2_i}}{\sum_i T_{1_i}} \geq 2 \\ \therefore f_{S(t)} &\leq 1/2 \end{aligned} \tag{17}$$

Thus, Equation 17 defines the necessary condition for a state to be stable in the ising formalism.

#### *9.2. Frustration and stability of k positive state in an n team network*

In a $n$-team network with no impurity, assuming uniform density ($d$) across all submatrices of the interaction matrix (within team and across team interactions) and equal team size ($T$) for all teams ($n * T$ nodes in total), the total number of edges is $d(nT)^2$. For a $k - positive$ state (i.e., $k$ teams have all nodes with expression 1 and $n - k$ teams have all nodes with expression $-1$). The interaction matrix can be divided into $n^2$ submatrices, with $n$ diagonal submatrices having $dT^2$ 1's and $(1 - d)T^2$ 0's. For $k - positive$ states, the edges in the diagonal submatrices ($dT^2$) will not be frustrated.

Let us now reorder the interaction matrix, keeping the $k$ teams that are on first, leading to four submatrices of the sizes :

1. $M1 : kT\ X\ kT$ corresponding to the teams that are on
2. $M2 : (n - k)T\ X\ (n - k)T$ corresponding to the teams that are off
3. $M12 : kT\ X\ (n - k)T$ edges from the $k$ teams to the $n - k$ teams
4. $M21 : (n - k)T\ X\ kT$ edges from the $n - k$ teams to the $k$ teams

The edges in $M12$ and $M21$ are not frustrated, since the edges in these matrices are all inhibitory, and the nodes participating in each edge have opposite expressions as well.

In $M1$ and $M2$, the diagonal elements are not frustrated, since the diagonals are all activating links (or don't exist), and the product of the expression of a node with itself is 1 (or 0). The non-diagonal elements will all be frustrated in both $M1$ and $M2$ and therefore are the only frustrated edges in the interaction matrix. Therefore,

$$\begin{aligned} frustration_{k-positive} &= \frac{d(kT)^2 - dkT^2 + d((n-k)T)^2 - d(n-k)T^2}{d(nT)^2} \\ &= \frac{k^2 + (n-k)^2 - n}{n^2} \\ &= \frac{n^2 - (2k+1)n + 2k^2}{n^2} \end{aligned}$$

Thus, following Equation 17, for a $k - positive$ state to be stable, the necessary condition to be met is

$$\begin{aligned} frustration_{k-positive} &\leq 0.5 \\ \frac{n^2 - (2k+1)n + 2k^2}{n^2} &\leq 0.5 \\ n^2 - (4k+2)n + 4k^2 &\leq 0 \end{aligned}$$

The above equations suggests the upper and lower limits for the number of teams for $k_{positive}$ states to be stable. Note that the quadratic function in $n$ presented above is concave upward. Thus, the function will take negative values between its roots. As the function takes positive value at $n = 0$ ($4k^2$)the roots will both be positive values, and can be calculated as $\frac{2k+1-\sqrt{16k+4}}{2}$ and $\frac{2k+1+\sqrt{16k+4}}{2}$. Therefore, the condition for a $k_{positive}$ state to be stable with n teams is:

$$\begin{aligned} max(\lceil \frac{2k+1-\sqrt{16k+4}}{2} \rceil, k) \leq n \leq \\ max(\lfloor \frac{2k+1+\sqrt{16k+4}}{2} \rfloor, k) \end{aligned} \tag{18}$$

The corresponding values for each $k$ are given in Table S1

| t | $n_max$ | $n_min$ |
|---|---|---|
| 1 | 3 | 1 |
| 2 | 5 | 2 |
| 3 | 7 | 3 |
| 4 | 8 | 4 |
| 5 | 10 | 5 |
| 6 | 11 | 6 |
| 7 | 12 | 7 |
| 8 | 14 | 8 |
| 9 | 15 | 9 |
| 10 | 16 | 10 |

Table S1: Caption

Restating the conditions in which the above results hold true:

- The network has $n$ mutually inhibiting teams
- Each team has an equal number of nodes
- A uniform density is maintained across the interaction matrix in each relevant submatrix
- No impurities. That is, the interactions within a team are either positive or absent. Similarly, the interactions across teams are negative or absent.

*9.3. Frustration and stability of k-positive states in different configurations*
